# A dynamic phosphoproteome reveals stage-specific regulation of cytokinesis in Leishmania

**DOI:** 10.64898/2026.11.09.765289

**Authors:** Y. Romina Nievas, Charlotte McNiven, Adam Dowle, Jeremy C. Mottram

**Author notes:** Corresponding authors &.

## Abstract

*Leishmania* parasites undergo extensive morphological and metabolic remodelling during their transition from motile insect promastigotes to non-motile intracellular amastigotes, yet how dynamic protein kinase signalling drives this process remains poorly understood. Here, we present a high-resolution temporal phosphoproteome of *L. mexicana* promastigote to axenic amastigote differentiation, profiling 16,495 phosphosites on 3,101 proteins across seven developmental time points and identifying 5,795 dynamically regulated sites. This temporal map reveals stage-specific signalling programs, including regulated sites in MPK2 and non-regulated sites in MRK1, protein kinases known to be essential for amastigote differentiation. Using analogue-sensitive (AS) chemical genetics, we show that acute inhibition of either kinase in axenic amastigotes disrupts the spatial coordination of cleavage furrow ingression with nuclear and kinetoplast segregation, revealing essential cytokinesis roles obscured in constitutive gene deletions. Despite generating similar cytokinesis phenotypes, genetic over-expression and cellular localisation analyses demonstrate that MPK2 and MRK1 act through spatially segregated signalling pathways. Time-resolved phosphoproteomics upon MPK2 inhibition places its activity upstream of cytoskeletal remodelling networks. Importantly, we show that the stage-specific requirement for MPK2 is defined by the parasite’s morphology: while motile promastigotes tolerate MPK2 inhibition, genetic ablation of flagellar motility sensitises them to loss of MPK2 activity. Thus, flagellar motility acts as a mechanical buffer during promastigote division, explaining why non-motile amastigotes are vulnerable to MPK2 inhibition. Together, our findings provide a systems level framework for *Leishmania* signalling and demonstrate that cellular architecture and biomechanics dictate protein kinase dependency across developmental stages of the parasite.

## INTRODUCTION

*Leishmania* are digenetic protozoan parasites that cycle between an extracellular stage within a phlebotomine sand fly vector and the intracellular niche in the mammalian host. Parasite survival and pathogenesis depend on successful adaptation to these contrasting environments. Promastigote-to-amastigote differentiation involves a profound alteration of parasite morphology, metabolism and cellular architecture, as highly motile, flagellated extracellular promastigotes transform into intracellular, rounded, non-motile amastigotes ^1,2^. Given that *Leishmania* lacks conventional gene-specific transcriptional control, post-transcriptional mechanisms and post-translational modifications (PTMs) play important roles in regulating cellular adaptation ^3^. In particular, protein phosphorylation provides a rapid, reversible mechanism to regulate protein function, subcellular localisation and complex assembly in response to developmental and environmental cues ^4^.

The *Leishmania* kinome is highly diverse and contains stage-essential protein kinases, indicating that developmental transitions are coordinated by distinct signalling programmes. Our previous high-throughput loss-of-fitness screening of protein kinase null mutants during *L. mexicana* promastigote-to-amastigote differentiation identified 29 and 43 protein kinases required for amastigote and promastigote survival, respectively ^5^. However, constitutive gene deletion strategies suffer from inherent limitations, as they cannot resolve the precise temporal onset of kinase activity during differentiation, distinguish roles in the initial differentiation process from subsequent amastigote maturation, or determine whether protein kinases required in promastigotes remain functionally important in amastigotes. Furthermore, while previous phosphoproteomic studies comparing defined developmental stages identified stage-specific phosphorylation signatures, endpoint comparisons of promastigote and amastigote stages fail to capture the temporal hierarchy of signalling events ^6–8^. Earlier time-course phosphoproteomic analysis provided initial evidence of dynamic phosphorylation changes during differentiation, but phosphosite coverage remained limited^9^. Consequently, the temporal organisation of phosphorylation events across differentiation, and how these dynamics relate to stage-specific kinase function, remains incompletely resolved.

In the mammalian host, successful infection demands not only differentiation but also sustained proliferation of the adapted amastigote stage. In *Leishmania*, cytokinesis requires precise spatiotemporal coordination of cleavage furrow ingression with the duplication and segregation of single-copy organelles, including the nucleus, kinetoplast and flagellar pocket ^10,11^. Given the substantial morphological divergence between life cycle stages, including the presence of an elongated motile flagellum in promastigotes versus a rudimentary, non-motile flagellum in amastigotes, the structural and biomechanical requirements for division may differ between stages.

In *Trypanosoma brucei*, flagellar structure and motility are closely linked to cytokinesis, where mechanical forces generated by the flagellum aid in furrow positioning and cell cleavage ^12,13^. Whether flagellar motility plays a similar structural or biomechanical role in *Leishmania* cell division remains unknown.

In this study, we applied quantitative, high-resolution phosphoproteomics across seven key time points during *L. mexicana* promastigote-to-axenic amastigote differentiation to map the dynamic phosphorylation landscape and identify stage-associated phosphorylation programmes. Among the protein kinases implicated in *Leishmania* differentiation, the MAP protein kinase MPK2 and the STE-family protein kinase MRK1 represent potential regulatory hubs ^5^. Here, we show that constitutive deletion of either kinase yields a striking convergent phenotype during differentiation: parasites initiate morphological adaptation and DNA replication but fail to undergo cytokinesis, accumulating as abnormal, multinucleated cells. This phenotypic similarity prompted us to investigate whether MPK2 and MRK1 function in a common signalling pathway controlling amastigote cytokinesis, whether their activities are strictly required during differentiation or for ongoing amastigote replication, and how they coordinate parasite division across different life-cycle stages.

Using chemical genetics, we demonstrate that MPK2 and MRK1 contribute independently to the spatial organisation of amastigote cytokinesis. Unexpectedly, we discover that the stronger dependency for MPK2 in axenic amastigotes is mechanically restricted, as while motile promastigotes tolerate MPK2 loss or inhibition, ablation of flagellar motility sensitises them to loss of MPK2 activity, demonstrating that flagellar beating serves as a mechanical safeguard during promastigote division. Finally, phosphoproteomic profiling following acute MPK2 inhibition identified downstream changes in cytoskeletal and cytokinesis-associated proteins. Together, our findings define the dynamic phosphoproteomic landscape during *Leishmania* differentiation and demonstrate that protein kinase dependency is dictated by an interplay between signalling networks, stage-specific cellular architecture and biomechanics.

## RESULTS

### Dynamic phosphoproteome reprogramming during differentiation

We recently identified 29 protein kinases essential for amastigote survival through a Bar-seq screen of protein kinase null mutants ^5^. To distinguish between kinases regulating the transient differentiation window and those operating in mature amastigote signalling, we performed a temporal phosphoproteomics analysis during *in vitro* promastigote to amastigote differentiation.

To capture the differentiation programme, samples of the *Leishmania mexicana* M379 cell line were collected from logarithmic procyclic promastigotes (LOG), stationary-phase promastigotes containing metacyclic promastigotes (STAT) and axenic amastigotes at 2 h (AxA2), 8 h (AxA8), 24 h (AxA24), 48 h (AxA48) and 5 days (AxA5D) after induction of metacyclic promastigote to amastigote differentiation with pH and temperature shift. Parasites retained a long flagellum through AxA2, which shortened at AxA8. From AxA24 onwards, the flagellum was no longer visible and the cells became rounder and smaller (Fig. 1a). In total, we identified 16,495 phosphosites present on 3,101 proteins, of which 5,795 sites on 1,888 proteins were significantly different during differentiation, revealing extensive highly coordinated temporal remodelling of the phosphoproteome (Table S1). Early differentiation time points showed a relatively modest number of changes (546, 459 and 310 different sites at STAT, AxA2 and AxA8 respectively) and were dominated by dephosphorylation. Phosphosite changes then expanded 5.3-fold between AxA8 and AxA24 (310 to 1,648 sites), rising to 2,257 at AxA48 and reaching a maximum of 4,159 sites in AxA5D, 72% of all sites that changed (Fig. 1b). This expansion was initially driven by increased phosphorylation (1,100 of 1,648 sites at AxA24), whereas mature amastigotes showed a net loss (2,371 of 4,159 sites decreased). Consistent with this, the hierarchical clustering of sample correlations resolved three groups: LOG, STAT, AxA2 and AxA8; AxA24 and AxA48; and AxA5D, placing the major transition between AxA8 and AxA24 (Fig. S1), the interval at which the flagellum is no longer visible (Fig. 1a).

To resolve the precise temporal dynamics of these events, we applied fuzzy C-means clustering, which identified 10 distinct phosphosite trajectories (Fig. 1c). For biological interpretation, clusters were further grouped into three higher-order classes according to the time point at which their average profile peaked: promastigote-associated (PRO; cluster 1-3, peaking at LOG), transition-associated (TRA; clusters 4-7, peaking at AxA24-AxA48) and amastigote-associated (AMA; clusters 8-10, peaking at AxA5D). The PRO clusters showed a progressive loss of phosphorylation upon differentiation. Cluster 1 decreased sharply during the LOG to STAT transition and was enriched in flagellar motility, cell cycle regulation, transcriptional repression and TOR signalling, consistent with reduced proliferation and increased motility in stationary-phase metacyclic promastigotes. Clusters 2 and 3 decayed mid-differentiation (AxA2-AxA8) and had a phosphorylation decrease in metabolic processes (purine and carbohydrate), flagellar motility and intraflagellar transport (IFT). These regulated processes mapped to the axoneme, IFT complexes and spindle structures, suggesting dismantling or down-regulation of motility and replicative machinery and restructuring of metabolic pathways (Fig. 1c-d).

TRA clusters comprised sites transiently phosphorylated during mid-differentiation. Cluster 4 captured the earliest transition events, rising continuously from LOG to AxA24 before declining, with enrichment in nutrient sensing, TOR signalling, chromatin remodelling and cell proliferation. Cluster 5 remained stable until AxA48, and dropped sharply at AxA5D, suggesting these phosphorylation states are transient and not maintained in mature amastigotes. This cluster was enriched in stress responses, transcription and translation control, and morphological changes (axoneme, cortical cytoskeleton and cell tip). Cluster 6 decreased in STAT, increased steadily through AxA24 and AxA48h, and declined again at AxA5D while Cluster 7 peaked at AxA24 and AxA48 and decreased at AxA5D. These clusters were enriched in cell cycle, transcription and translation control and TOR/ARF signalling terms, mapping to cellular components such as the basal body, flagellar pocket and cell tip. Collectively, these profiles suggest TRA clusters coordinate nutrient sensing, cell cycle progression and transcriptional reprogramming during the differentiation transition (Fig. 1c-d).

AMA clusters showed increased phosphorylation at late time points. Clusters 8 and 9 peaked at AxA24. Cluster 9 maintained elevated levels through AxA48 and AxA5D and was enriched in post-transcriptional regulation, pyrimidine biosynthesis and heat response, mapping to the basal body, cell tip, cortical cytoskeleton and the chromatin remodelling ISWI complex. Cluster 8 displayed a biphasic profile, peaking at AxA24, stabilising at AxA48 and peaking again at AxA5D. Terms were enriched in cytoskeleton organisation, mRNA capping and splicing, glutamine and phospholipid metabolism, and TOR signalling, with cellular components spanning the flagellar pocket, acidocalcisome and the mitochondrial outer membrane. Cluster 10 exhibited a late peak at AxA5D, enriched in cytokinesis, translation regulation and phosphoinositide (PIP)/TOR signalling, localising to the flagellar pocket, cell tip and nuclear body. Together, these AMA clusters coordinate membrane, cytoskeleton and organelle remodelling, alongside the post-transcriptional and metabolic adaptations characteristic of the amastigote stage (Fig. 1c-d).

To assess phosphosite dynamics in the context of global proteome changes, we compared temporal protein abundance profiles with their corresponding phosphosite profiles while preserving the phosphoproteomic clustering structure (Fig. 1c). Overall, phosphorylation changes showed weak correlation with protein abundances (r = 0.13 to 0.41) and only 22-31% of the parent proteins themselves were significantly modulated, indicating that phosphorylation dynamics are largely independent of changes in protein abundance. The exceptions were the three PRO clusters, characterized by decreasing phosphorylation, where the correlation was moderate (mean r = 0.41-0.56) and 47 to 60% of parent proteins were also modulated. Notably, the decrease in PRO cluster phosphorylation was temporally accompanied by an increase in phosphorylation of proteins involved in ubiquitination/deubiquitination and dephosphorylation processes (detected in clusters 4, 8, 9 and 10), suggesting coordinated activation of post-translational regulatory networks during differentiation. Volcano plots highlighting each contrast and significant phosphosites are provided in Fig. S2 and Table S1.

**Figure 1.**
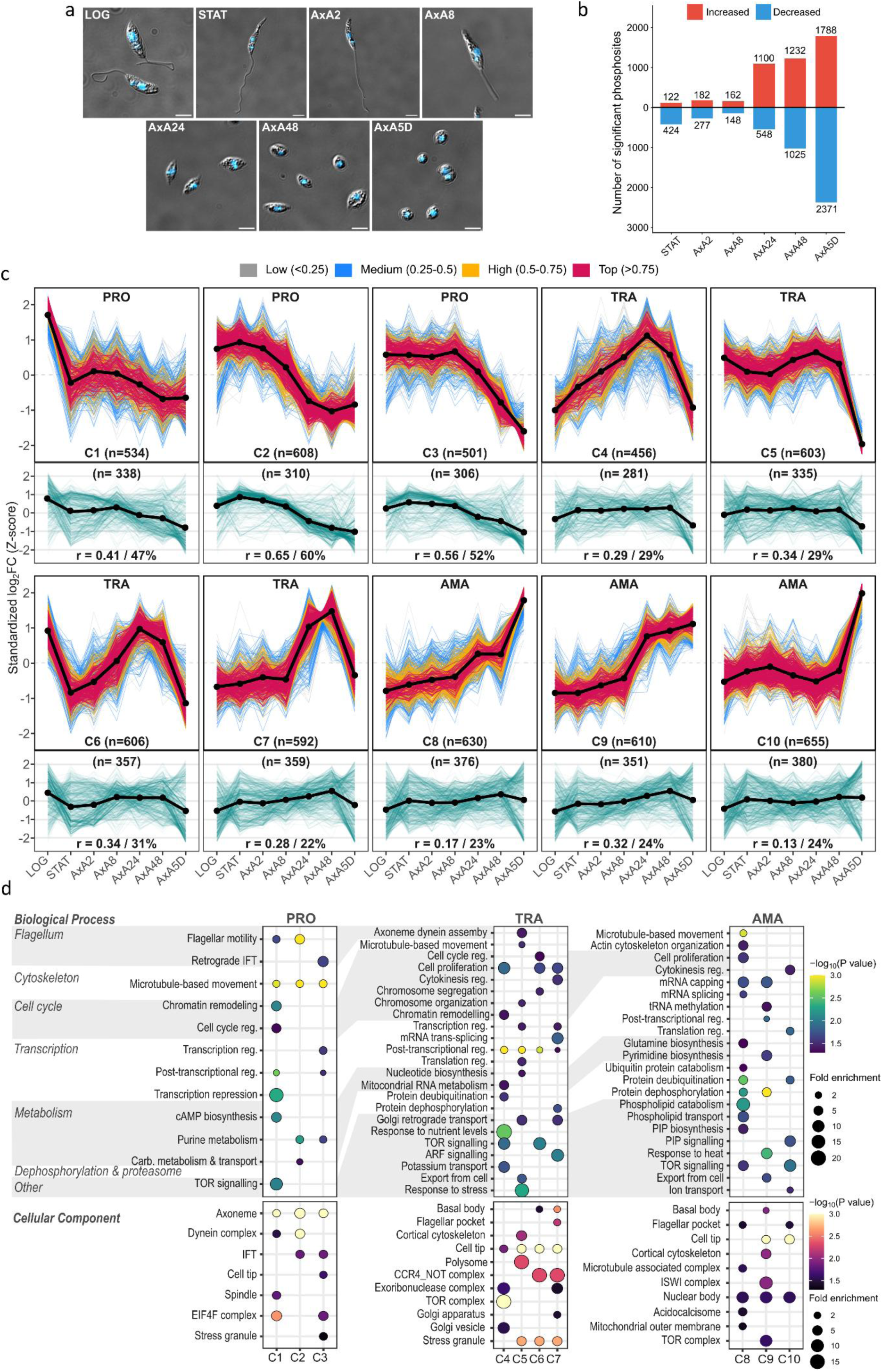
Global phosphoproteomic dynamics during *L. mexicana* differentiation. (a) Representative micrographs of *L. mexicana* at each time point of the differentiation time course. DNA was stained with Hoechst (blue). Scale bar: 2 µm. (b) Absolute quantification of significantly increased (red) or decreased (blue) phosphosite abundance across the indicated differentiation time points. (c) Mfuzz clustering of significantly modulated phosphosites (multicolour lines; membership values indicated; black lines are cluster centroids) aligned with the abundance profiles of their corresponding parent proteins (teal lines below; black lines are arithmetic means). Total sites/proteins per cluster are indicated (n). Pearson correlation coefficients (r) assess the kinetic trajectory coupling between phosphosites and total proteins. The percentage of those proteins that are themselves significantly modulated in abundance is also indicated. Axes denote standardised log_2_ fold changes (Y-axis) across the experimental time course (X-axis). LOG, logarithmic procyclic promastigotes; STAT, stationary-phase promastigotes containing metacyclic promastigotes; AxA2, AxA8, AxA24, AxA 48 and AxA5D, axenic amastigotes at 2 h, 8 h, 24 h, 48 h and 5 days after differentiation. PRO (Promastigote); TRA (Transition); AMA (Amastigote). C1 to C10 indicate Clusters 1 through 10, respectively. (d) Biological process and cellular component gene ontology terms enriched in the different clusters.

### Dynamic phosphorylation of protein kinases required for amastigote and promastigote survival

Having established the global phosphoproteomics landscape of differentiation, we next examined phosphorylation dynamics across the 29 protein kinases previously identified as essential for amastigote survival. We detected significantly regulated phosphosites in 22 of them, predominantly associated with the TRA and AMA phosphorylation programmes. Based on the temporal distribution of regulated phosphosites, protein kinases could be stratified into three classes; those regulated early in differentiation, those displaying phosphorylation changes throughout both differentiation and amastigote maturation, and those predominantly regulated in mature amastigotes.

Several protein kinases had phosphorylation changes primarily during the transition phase (Fig. 2). The nutrient and osmotic stress sensor TOR3 ^14^ contained three regulated phosphosites, located between the kinase and FATC domains. Similarly, the calcium/calmodulin kinase LmxM.32.1710 harboured a cluster of six regulated phosphosites within the linker region separating the kinase domain from the first EF-hand calcium binding motif, most of these displayed increased phosphorylation early during differentiation (STAT). The amastigote-stage regulated MPK10 ^15^ displayed transient phosphorylation, with both detected sites peaking at AxA24 and declining thereafter. Additionally, protein kinase LmxM.06.0640, displayed five sites modulated during the transition, located mainly within the large N-terminal disordered region (Fig. 2).

A second group of protein kinases displayed phosphorylation changes spanning both differentiation and amastigote maturation. These included CK2A1, CK2A2, TTBK, MPK15, EIF2AK2 and protein kinase LmxM.07.0250, all of which contained phosphosites assigned to both TRA and AMA phosphorylation programmes, indicating multilayered post-translational regulation across developmental progression.

In contrast, a distinct subset of protein kinases were regulated predominantly during amastigote maturation. AKB1, implicated in cytokinesis in *T. brucei* ^16^ contained three AMA-associated phosphosites distributed across both the terminal region and the kinase domain. ZFK, another differentiation-associated kinase ^17^ also displayed multiple AMA-enriched phosphosites. MPK1, essential for *Leishmania* amastigotes ^18^, displayed two AMA-regulated sites between its DFG and APE variant (SPE) motifs. MPK2, previously linked to arginine sensing and antimony resistance, contained three regulated phosphosites within its kinase domain. These included the two largest phosphorylation increases detected among all amastigote essential kinases, both located within the canonical TDY activation motif (Log_2_FC^T174^ = 5.13; Log_2_FC^Y176^ = 4.98) and previously identified as phosphorylation sites ^19,20^. Additionally AMA-regulated protein kinases included LmxM.25.1560, LmxM.06.1180, LmxM.16.0300, LmxM.21.0270 and LmxM.34.2870.

To determine whether signalling components important for the promastigote stage also undergo phosphorylation during differentiation, we extended this analysis to the 43 protein kinases previously identified as required for promastigote fitness ^5^. Twenty of these displayed significant phosphoregulation during differentiation (Fig. S3), spanning PRO, TRA and AMA programmes. Among these, DYRK1, implicated in stationary phase maintenance ^21^ and PKAC1, involved in flagellar motility ^22^, showed phosphoregulated sites across PRO and TRA programmes. TOR1, involved in nutrient-controlled proliferation ^23^, displayed regulated sites mainly during the transition, whereas TOR2 contained two sites regulated in the transition and six additional sites with increased phosphorylation in AMA (AxA5D). CK1.2, a protein kinase released into the macrophage and implicated in host-pathogens interactions ^24^, exhibited a regulated site during the transition within the ATP-binding region. Cell cycle associated kinases, including CRK12 ^25^, CRK9 and KKT3 ^26^ showed regulated phosphosites during the TRA programme, whereas CRK3 ^27^, GSK3 ^28^ and GSKA were predominantly regulated in the AMA programme. CLK1 and CLK2 exhibited distinct phosphorylation dynamics, with uniquely identifiable phosphopeptides from CLK1 and CLK2 showing divergent profiles. CLK1 displayed an oscillatory profile consistent with cell cycle-associated regulation, whereas CLK2 phosphorylation remained elevated in mature amastigotes. Thus, a substantial proportion of protein kinases required for promastigote viability are actively and dynamically rewired during the differentiation process. Across both amastigote and promastigote essential protein kinases, the majority of regulated phosphosites are located outside the kinase domain (90 sites outside versus 41 sites within), clustering predominantly within predicted disordered regions flanking the catalytic domain (Fig. 2, Fig. S3).

**Figure 2.**
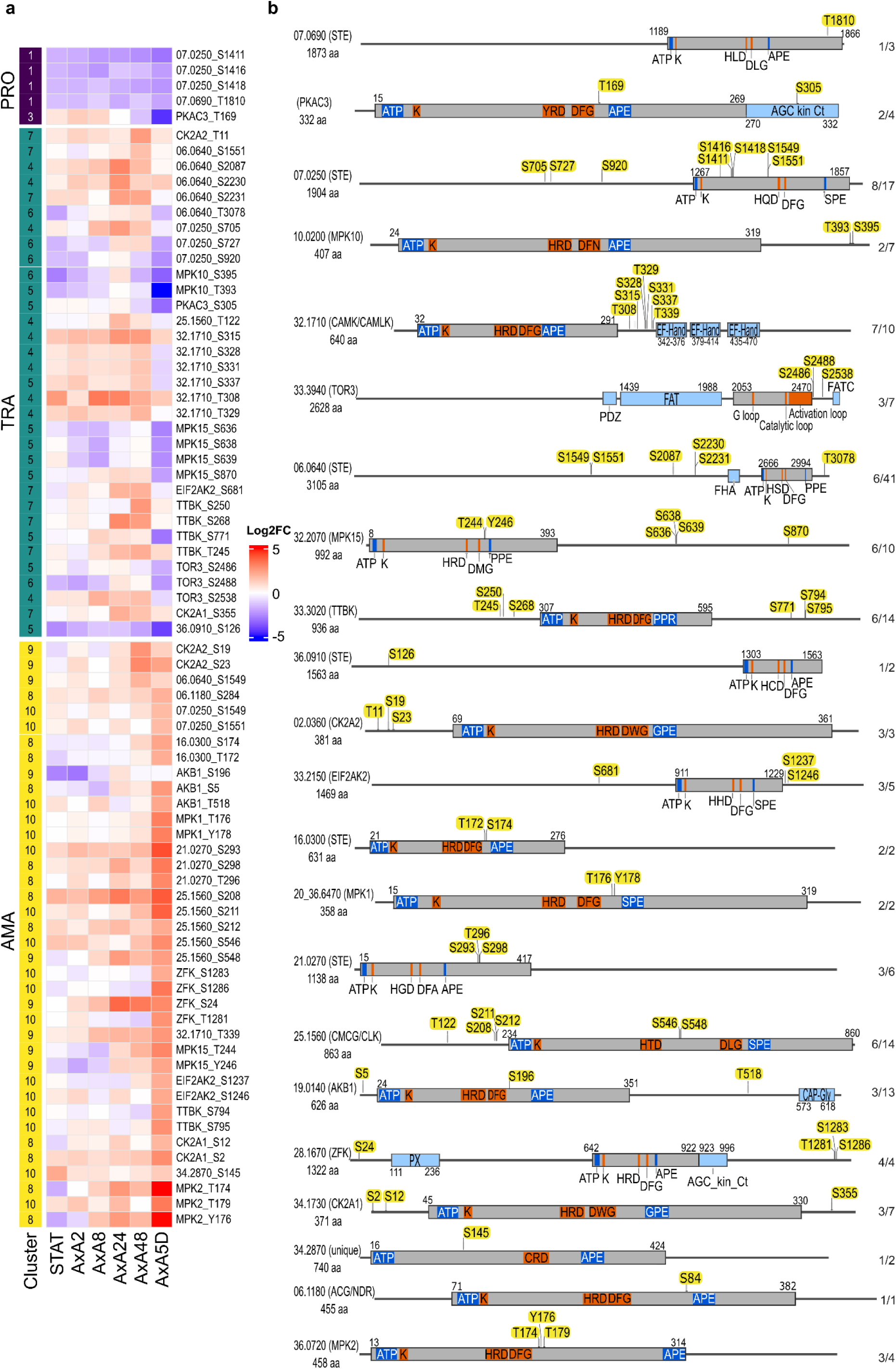
Structural architecture and phosphorylation dynamics of essential protein kinases during differentiation. (a) Heatmap displaying Log_2_ fold-change values of modulated phosphosites across distinct developmental stages: Promastigote PRO (Promastigote); TRA (Transition); AMA (Amastigote). Phosphosites were partitioned into categories derived from Mfuzz clustering. LOG, logarithmic procyclic promastigotes; STAT, stationary-phase promastigotes containing metacyclic promastigotes; AxA2, AxA8, AxA24, AxA 48 and AxA5D, axenic amastigotes at 2 h, 8 h, 24 h, 48 h and 5 days after differentiation. (b) Schematic representation of the 22 identified protein kinases with modulated phosphorylation sites (indicated by yellow markers). Core kinase domains are depicted as gray bars, indicating key catalytic landmarks: the ATP-binding site (ATP; blue), the catalytic lysine (K; orange), the catalytic loop aspartate within the HRD motif and its variants (YRD, HTD, HCD, HGD, HQD, HLD, HSD; orange), the metal-coordination aspartate within the DFG motif and its variants (DLG, DFA, DFN, DMG, DWG; orange) and the activation loop anchor APE motif and its variants (GPE, SPE, PPE; blue). Additional structural and regulatory domains are represented by light blue bars, including Forkhead-associated (FHA), Phox (PX), AGC kinases C-term regulatory (AGC kin Ct), calcium binding EF hand (EF-Hand), Cytoskeleton Associated Protein Glycine-rich (CAP-Gly), FAT/FATC regulatory and PDZ scaffolding domains. Domain boundaries were identified using the PROSITE database. Core catalytic motifs were defined using multiple sequence alignment with MAFFT-SparseCore G-INS-i global iterative method. Sequence maps were generated using SnapGene software (Dotmatics). Numbers besides each diagram indicate modulated/total detected phosphosites. Gene IDs have been shortened for clarity (eg LmxM.07.069 to 07.0690). Protein kinase names or families are indicated in parentheses, followed by protein length in amino acids (aa).

### MPK2 and MRK1 null mutants exhibit a cell division defect during amastigote differentiation

We selected MPK2 and MRK1 for in-depth functional characterisation to resolve how distinct phosphorylation profiles map to stage-specific essentiality. MPK2 emerged from our phosphoproteomic analysis as a prime candidate regulator of amastigote biology, displaying the highest magnitude phosphorylation increase in mature amastigotes, whereas MRK1 maintained a static phosphorylation profile throughout differentiation. Surprisingly, despite these opposing dynamics, Δ*mpk2* and Δ*mrk1* parasites exhibited very similar cytokinesis arrest during differentiation (Fig. 3). Investigating both protein kinases allowed us to address whether dynamic phosphoregulation is required to execute amastigote division, and whether these phenotypically convergent protein kinases function in a common pathway or parallel cascades.

We first assessed the individual fitness of the Δ*mpk2* and Δ*mrk1* cell lines during differentiation to axenic amastigotes. Complemented lines were generated by integrating each gene into the ribosomal locus of the respective null mutants (Δ*mpk2*::*MPK2* and Δ*mrk1*::*MRK1*). Viability assays performed 5 days after induction of differentiation, revealed that Δ*mrk1* and Δ*mpk2* lines exhibited a 70% and 90 % reduction in viability, respectively, compared to the parental control (T7Cas9) (Fig. 3a). Importantly, genetic complementation restored viability to 90% for Δ*mrk1*::*MRK1* and 70% for Δ*mpk2*::*MPK2* after 5 days of differentiation, indicating that the observed loss-of-fitness (LOF) phenotypes are attributable to the absence of these protein kinases (Fig. 3a).

Because differentiation requires coordinated cell-cycle progression and extensive morphological remodelling, including flagellar retraction and cell body rounding, we next monitored cell-cycle progression and morphology during this transition. In stationary-phase metacyclic promastigotes, both null mutants displayed a cell cycle profile similar to T7Cas9, with the majority of the cells in G1, as expected (Fig. 3b). However, 24 h after differentiation, both Δ*mrk1* and Δ*mpk2* showed an accumulation of cells in the G2 phase, followed by the appearance of multinucleated cells after 72 h (Fig. 3b). Although both null mutants completed flagellar retraction by 72 h (Fig. 3c), Δ*mrk1* cells were significantly longer and wider, while Δ*mpk2* cells were slightly shorter but wider than T7Cas9 (Fig. 3d). The presence of bi-nucleated and multinucleated cells observed by microscopy was consistent with the G2-arrest phenotype identified by flow cytometry.

We next evaluated the expression dynamics and subcellular distribution of MRK1 and MPK2. Protein abundance across differentiation was assessed from the total proteome dataset. MRK1 abundance did not change significantly at any time point relative to LOG, indicating MRK1 protein levels remain stable during differentiation. MPK2 abundance likewise remained stable through 48 h post induction, but increased approximately two-fold in mature axenic amastigotes (AxA5D) (Fig. 3e). This indicates that during early differentiation MPK2 activity is modulated primarily through phosphorylation, while the increase in mature amastigotes points to an additional, later contribution from elevated protein levels. (Fig. 3e). Regarding localisation, MRK1 accumulated at the posterior pole in stationary phase promastigotes and became cytoplasmic 24 h post-differentiation. Conversely, MPK2 consistently accumulated at both the anterior and posterior tips in both stationary phase promastigotes and 24 h differentiated amastigotes (Fig. 3f).

Collectively, these results show that while Δ*mrk1* and Δ*mpk2* cells can initiate morphological differentiation and replicate their DNA, they are blocked in their ability to complete cell division.

**Figure 3.**
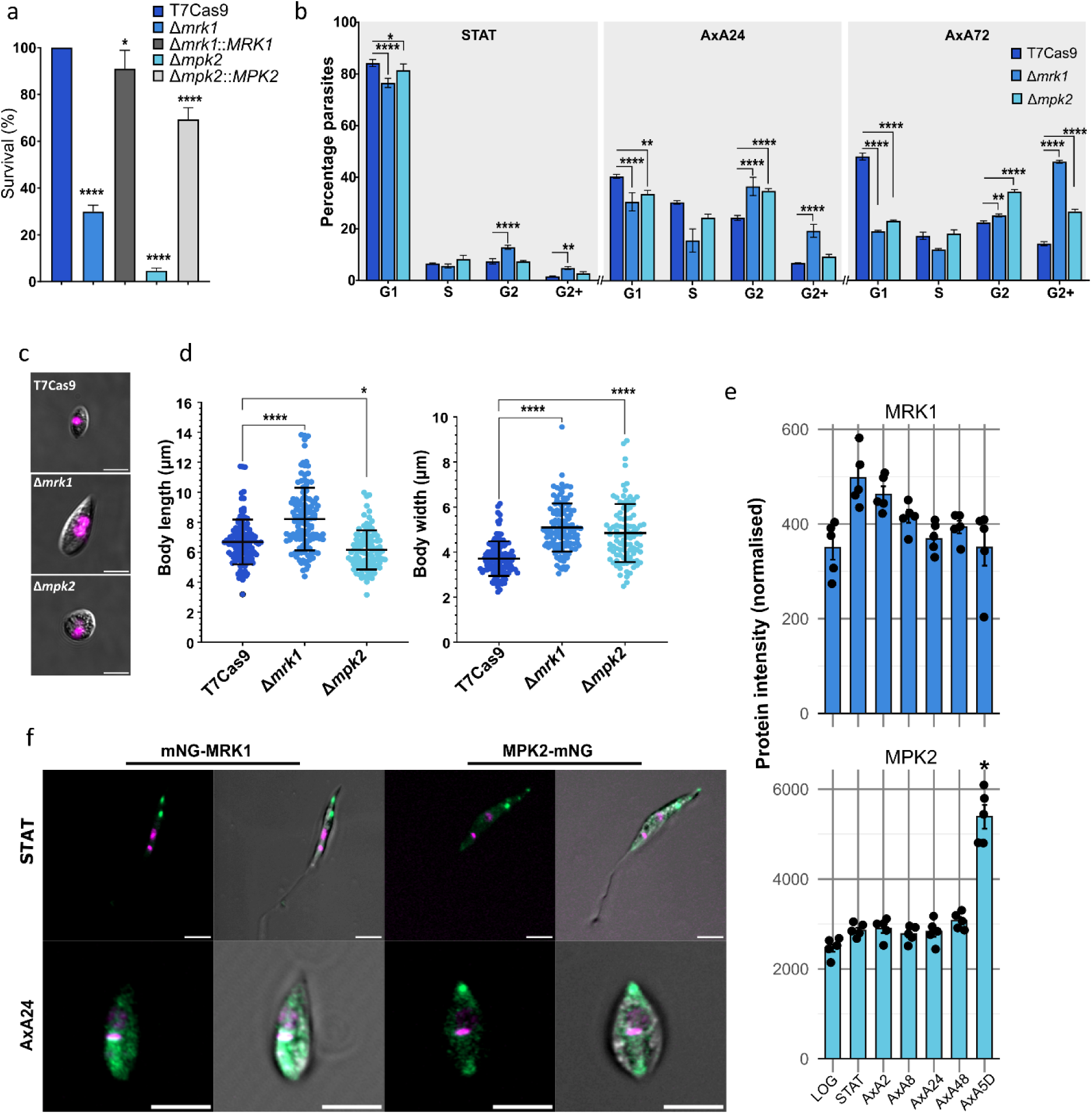
MRK1 and MPK2 are essential for promastigote to amastigote differentiation. **(a)** Viability of protein kinase null mutants (Δ*mpk2* and Δ*mrk1)* and complemented lines (Δ*mpk2*::*MPK2* and Δ*mrk1*::*MRK1*) assessed by Alamar Blue assay 5 d after induction of differentiation, normalised to T7Cas9 (set at 100%); significance assessed relative to T7Cas9 by ordinary one-way ANOVA with Dunnett’s multiple comparison test. (b) Cell-cycle progression during differentiation, monitored by propidium iodide staining and flow cytometry. The percentages of cells in G_1_, S and G_2_ and those with DNA content greater than G_2_ (G_2_+) are reported. Comparisons were made within each cell-cycle phase (G_1_, S, G_2_, G_2_+) and significance was assessed relative to T7Cas9 by two-way ANOVA with Dunnett’s multiple comparison test. (c) Representative micrographs after 3 days of differentiation. DNA was counterstained with Hoechst (magenta). Scale bar: 5 µm (d) Morphometric analysis of cell body width and length (individual cells from two independent biological replicates were pooled and measured using ImageJ); significance assessed relative to T7Cas9 by ordinary one-way ANOVA with Dunnett’s multiple comparison test. (e) MRK1 and MPK2 protein abundance across differentiation, from the total proteome dataset (bars show mean normalised intensity ± SEM; points show individual biological samples); significance assessed relative to LOG by two-sided moderated t-test (limma, via DEP2) with Benjamini–Hochberg FDR correction (asterisks in e indicates *Padj* < 0.05 with log2FC ≥ 1). (f) Subcellular localisation of endogenously N-terminally mNeonGreen-tagged MRK1 and C-terminally mNeonGreen-tagged MPK2 (green) visualised by live-cell fluorescence microscopy in STAT and AxA24 cells. DNA was counterstained with Hoechst (magenta). Scale bars: 5 µm. Data represent mean ± SD of three biological replicates unless otherwise indicated, exact n, P values and individual data points are provided in the Source Data file. *P < 0.05;**P < 0.01; ***P < 0.001, ****P < 0.0001.

### MPK2 and MRK1 kinase activity is required for amastigote differentiation and replication

Because the lethal loss-of-function phenotype of MPK2 and MRK1 null mutants prevented us from studying their roles in fully differentiated amastigotes, we established a chemical genetics approach to enable acute protein kinase chemical inhibition across distinct life-cycle stages. Using precision editing, we mutated the gatekeeper residue to alanine to generate analogue-sensitive (AS) lines, creating an expanded ATP-binding pocket that accommodates bulky kinase inhibitors (BKIs), such as PP1-derived analogues. As controls, we introduced a synonymous mutation at the same locus to generate silent, inhibitor-insensitive control lines ^26,29^. Mutations were confirmed by PCR and sequencing (Fig. S4).

We next evaluated the requirement for MPK2 and MRK1 kinase activities during differentiation by generating dose-response curves using the bulky ATP analogue 1NA-PP1. Both AS mutants showed high sensitivity to 1NA-PP1, displaying IC_50_ values of 0.4 µM for MPK2^F91A^ and 3.67 µM for MRK1^M392A^, compared to 3.93 µM for MPK2^F91F^ and 10.5 µM for MRK1^M392M^ silent control lines (Fig. 4a).

To test if protein kinase activity is also required after differentiation is complete, mature amastigotes were treated with 1NA-PP1. Both AS mutants remained significantly more sensitive to inhibition than their respective controls, displaying IC_50_ values of 3.49 µM for MPK2^F91A^ and 6 µM for MRK1^M392A^, compared to 22.1 µM for MPK2^F91F^ and 25.3 µM for MRK1^M392M^ (Fig. 4b).

We then investigated whether acute chemical inhibition recapitulated the null mutant phenotypes. We evaluated cell cycle progression in mature amastigotes following treatment with 15 µM 1NA-PP1, a concentration that maintained high viability in control lines (MPK2^F91F^, 77% and MRK1^M392M^, 88%) while severely reducing viability in the AS mutants (MPK2^F91A^, 9% and MRK1^M392A^, 2%). Prior to treatment, the AS and silent mutant lines displayed comparable cell-cycle profiles characteristic of proliferating amastigotes (0 h). After 12 h of inhibition, both MPK2^F91A^ and MRK1^M392A^ amastigotes treated with 1NA-PP1 displayed a marked increase in G_2_ phase cells compared to the untreated AS mutants and 1NA-PP1 treated silent mutant controls. By 24 h of treatment, the AS mutant phenotype became more pronounced, with the accumulation of multinucleated cells (G_2_+) mimicking the cytokinesis block observed with the null mutants (Fig 4c-d). These findings confirm that the catalytic activities of MPK2 and MRK1 are required for both promastigote-to-amastigote differentiation and mature amastigote replication.

**Figure 4:**
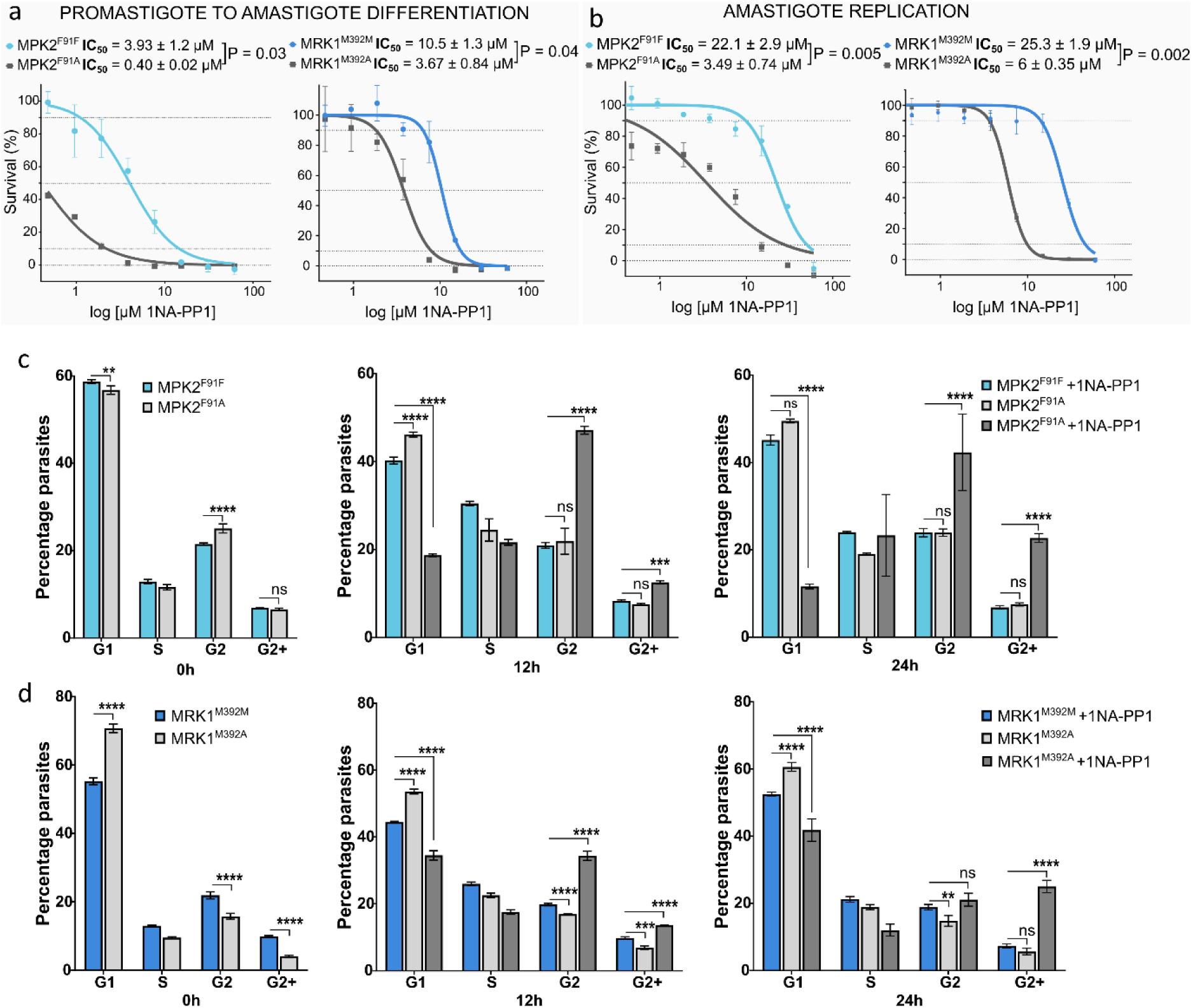
MPK2 and MRK1 protein kinase activity is required for promastigote to amastigote differentiation and for amastigote replication. Susceptibility of the AS mutants (MPK2^F91A^ and MRK1^M392A^) relative to their respective silent controls (MPK2^F91F^ and MRK1^M392M^), to 1NA-PP1 during (a) promastigote-to-amastigote differentiation and (b) mature amastigote replication. IC_50_ (mean ± SD) and exact *P* values are shown for each comparison (unpaired two-tailed t-test with Welch’s correction). (c-d) Cell-cycle progression of MPK2^F91F^/MPK2^F91A^ (c) and MRK1^M392M^/ MRK1^M392A^ (d) parasites, monitored by propidium iodide staining and flow cytometry, before 1NA-PP1 treatment (0 h), 12 h and 24 h after treatment. At 0 h, the AS mutant was compared to its untreated silent control; at 12 h and 24 h, the untreated and 1NA-PP1-treated AS mutant were each compared to the treated silent control. The percentages of cells in G_1_, S and G_2_ and those with DNA content greater than G_2_ (G_2_+) are reported. Comparisons were made within each cell-cycle phase (G_1_, S, G_2_, G_2_+) by two-way ANOVA on % of parasites per phase, followed by Sidak’s (0 h) or Dunnett’s (12 h / 24 h) multiple comparisons tests. Data represents mean ± SD of three biological replicates unless otherwise indicated. *P < 0.05; **P < 0.01; ***P < 0.001, ****P < 0.0001. Exact P values for (c, d) are provided in the Source Data file.

### MPK2 and MRK1 activities are required for spatial coordination of amastigote cytokinesis

To define the mechanistic roles of MPK2 and MRK1 during amastigote cytokinesis, we endogenously myc-tagged the kinases containing the silent and AS mutations, and characterised their subcellular localisation. MPK2 (C-terminal; MPK2^F91F^::myc) exhibited a dynamic distribution, as in both dividing (2N2K) and non-dividing (1N1K) cells it localised either at the anterior, posterior or both cell poles (Fig. 5b, e). During cytokinesis, MPK2^F91F^::myc was enriched within the cytokinetic bridge connecting the daughter cells and concentrated at the tips of β-tubulin positive microtubule bundles at both cell poles, suggesting an association with cytoskeletal structures undergoing remodelling during cell division (Fig. 5b-c). In contrast, MRK1 (N-terminal; myc::MRK1^M392M^) exhibited a predominantly diffuse cytoplasmic localisation in both non-dividing (1N1K) and dividing amastigotes (2N2K), indicating that while both kinases are essential for cytokinesis, they are spatially segregated (Fig. 5d).

To determine if kinase activity is required to maintain their subcellular organisation, MPK2^F91A^::myc and myc::MRK1^M392A^ mutants were treated with 1NA-PP1 and analysed by immunofluorescence. Inhibition of MPK2^F91A^::myc cells resulted in the loss of polarised MPK2 localisation pattern, with the protein kinase becoming diffusely cytoplasmic in both non-dividing and dividing cells. However, MPK2 remained detectable at the centre of the cytokinetic bridge in late cytokinesis. myc::MRK1^M392A^ localisation remained largely unaffected by inhibition, consistent with its diffuse cytoplasmic distribution (Fig. 5e-f).

We next investigated whether protein kinase inhibition altered the directional trajectory of the cleavage furrow. Cytokinesis in *Leishmania* procyclic promastigotes typically proceeds via an anterior-to-posterior cleavage furrow ingression, accompanied by daughter cell twisting and posterior abscission ^10,30^ . However, alternative furrowing trajectories, such as bidirectional, internal or posterior-to-anterior furrow progression, have also been reported ^11^, although prevalence and regulation in amastigotes remains unknown. In untreated T7Cas9 amastigotes, anterior-to-posterior furrow progression was the predominant trajectory (58.4 %), followed by bidirectional furrowing (28.6 %), and posterior-to-anterior furrowing (13 %) (Fig. 5g-h). Treatment of silent mutant controls with 1NA-PP1 did not significantly alter these distributions. In contrast, inhibition of MPK2^F91A^::myc and myc::MRK1^M392A^ resulted in a marked increase in posterior-to-anterior furrowing without affecting the frequency of bidirectional furrowing (Fig. 5h).

Because alternative furrowing can reflect a defect in coordinating cytokinesis with organelle segregation, we next quantified nuclear/kinetoplast (N/K) configurations associated with different furrow trajectories. In T7Cas9 amastigotes, posterior-to-anterior cytokinesis was correlated with a higher frequency of abnormal N/K configurations compared with anterior-to-posterior and bidirectional furrowing (Fig. 5i). Consistently, inhibition of MPK2^F91A^::myc and myc::MRK1^M392A^ significantly increased the frequency of abnormal N/K configurations compared with silent mutant controls. These abnormalities included 1K1N cells initiating furrow ingression prior to completing organelle segregation, as well as aberrant 1N2K and 2N1K configurations (Fig. 5f, j). Importantly, inhibited cells undergoing posterior-to-anterior furrowing were still capable of generating normal 2K2N daughter cells, indicating that organelle segregation can still proceed despite kinase inhibition (Fig. 5f).

Together, these results indicate that MPK2 and MRK1 kinase activities are required for the spatial organisation of amastigote cytokinesis. Disruption of their activity redirects cleavage furrow ingression toward an alternative posterior-to-anterior trajectory and increases division errors, suggesting that these protein kinases contribute to the temporal coordination of cytokinesis with nuclear and kinetoplast inheritance.

**Figure 5.**
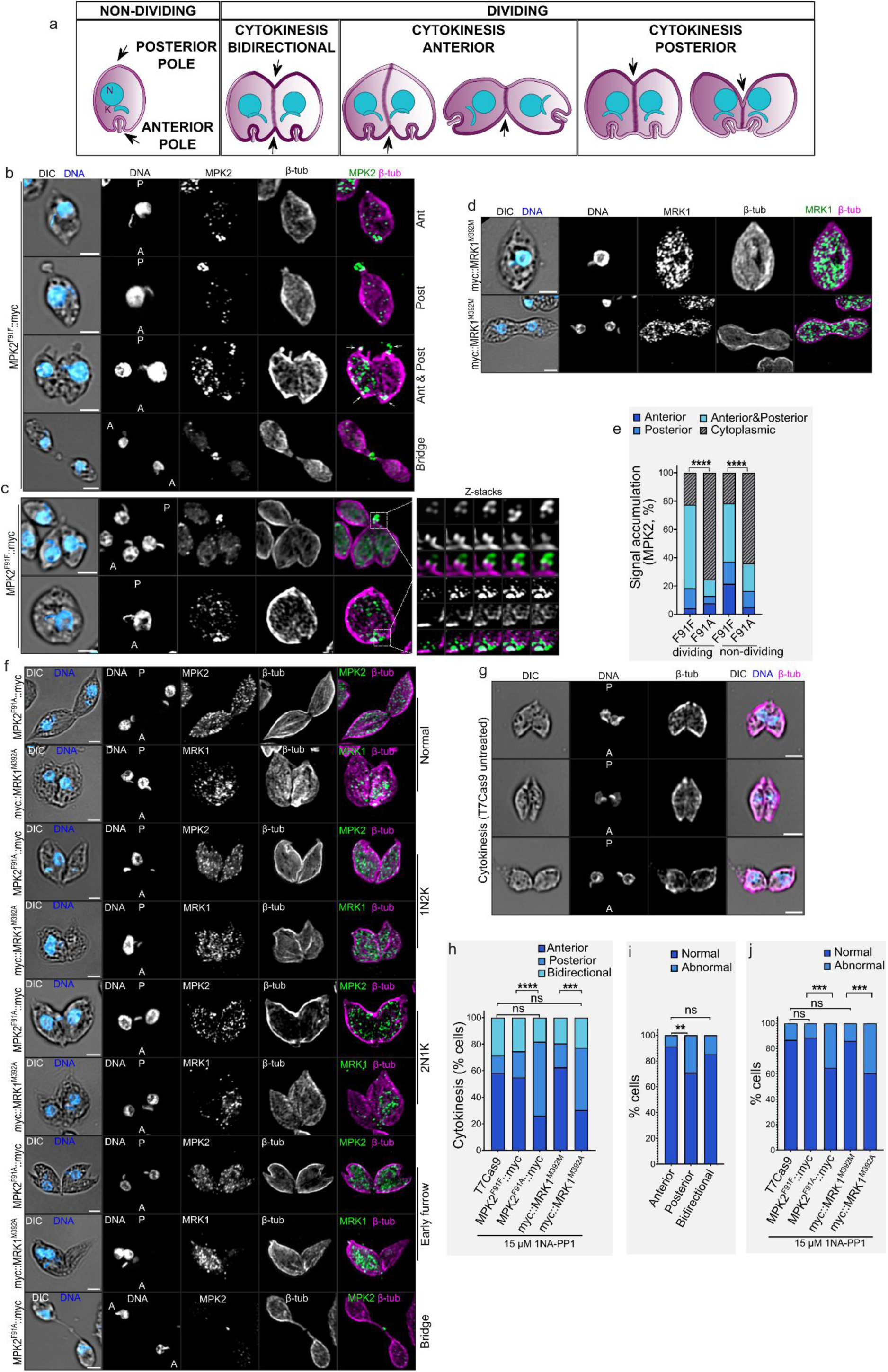
Localisation of MPK2 and MRK1 and phenotypic consequences of their inhibition during amastigote cytokinesis. Myc-tagged MPK2^F91F^, MRK1^M392M^ , MPK2^F91A^ and MRK1^M392A^ lines were treated with 15 µM 1NA-PP1 for 12 h followed by immunofluorescence. Protein kinases were labelled with anti-myc (green), microtubules with anti-β-tubulin (magenta) and DNA was counterstained with DAPI (blue). Scale bars: 2µm. A: anterior; P: posterior. (a) Schematic representation of parasite polarity and cytokinesis furrow progression. Nucleus (N); Kinetoplast (K). (b) Subcellular localisation of MPK2 in MPK2^F91F^::myc amastigotes. Representative cells show MPK2 enrichment at the anterior pole (Ant), posterior pole (Post), both poles (Ant & Post), or within the cytokinetic bridge (bridge). (c) Partial colocalisation of MPK2 with microtubule bundles at cell poles in MPK2^F91F^::myc parasites. Insets depict individual Z-stack sections. (d) Subcellular localisation of MRK1 in myc::MRK1^M392M^ cells. (e) Quantified distribution of MPK2 localisation patterns in dividing and non-dividing MPK2^F91F^::myc and MPK2^F91A^::myc cells. (f) Representative micrographs of myc::MRK1^M392A^ and MPK2^F91A^::myc lines undergoing posterior-to-anterior cytokinesis, illustrating normal (2N2K) and abnormal (1N2K, 2N1K, early furrowing) N/K configurations. (g) Micrographs of untreated T7Cas9 cells showing representative cleavage furrow progression trajectories. (h) Quantification of cleavage furrow trajectories across T7Cas9 and 1NA-PP1-treated MRK1 and MPK2 silent and AS lines. (i-j) Quantification of N/K distribution patterns. Cells were scored as *normal* when displaying two fully segregated nuclei and two kinetoplasts (2K2N) correctly positioned along the cellular axes. Cells were categorised as *abnormal* if they exhibited at least one of the following defects: incomplete nuclear or kinetoplast segregation, aberrant copy number (such as 1N2K, 2N1K, multinucleate, or multi kinetoplast), or mislocalisation of nuclear or kinetoplast DNA within the cell body; (i) Frequency in untreated T7Cas9; (j) Frequency in 1NA-PP1 treated MRK1 and MPK2 silent and AS lines. Data are presented as percentages; statistical significance was assessed using Fisher’s Exact Test on raw counts (sample sizes *n* and exact P values detailed in Source Data). *P < 0.05; **P < 0.01; ***P < 0.001, ****P < 0.0001.

### MRK1 and MPK2 act independently to regulate amastigote cytokinesis

The similar cytokinesis defects observed upon MPK2 and MRK1 inhibition suggested that these two protein kinases may operate within the same linear signalling pathway. To test this hypothesis, we episomally overexpressed each protein kinase in the silent or the AS mutant background of the reciprocal protein kinase, asking whether increased dosage of one protein kinase could compensate for the loss of activity of the other. Overexpression of both MPK2 and MRK1 was confirmed by western blot (Fig. S5).

We first assessed whether protein kinase overexpression rescued parasite viability during amastigote replication under 1NA-PP1 treatment. Overexpression of MPK2 in the MRK1^M392M^ control (MRK1^M392M^[MPK2]) did not affect viability. However, elevated levels of MPK2 during MRK1^M392A^ inhibition (MRK1^M392A^[MPK2]) led to a partial phenotypic rescue, resulting in a significant three-fold increase in viability compared to inhibited MRK1^M392A^ without over-expression (Fig. 6a). To investigate whether this rescue was due to restoration of normal cell division, we analysed cell-cycle progression by flow cytometry. Unexpectedly, MPK2 overexpression did not restore a normal cell-cycle profile of MRK1-inhibited parasites. Instead, it further increased the percentage of multinucleated cells (Fig. 6b). Conversely, overexpression of MRK1 in MPK2^F91F^ or MPK2^F91A^ (MPK2^F91F^[MRK1], MPK2^F91A^[MRK1]) failed to rescue cell viability or alter cell-cycle progression following 1NA-PP1 treatment (Fig. 6a, c).

Because shared subcellular localisation can provide evidence for functional coupling between proteins, we next investigated whether MPK2 and MRK1 occupied overlapping microdomains during cytokinesis. We generated endogenously double-tagged cell lines expressing MPK2::HA in MRK1 backgrounds (myc::MRK1^M392M^ and myc::MRK1^M392A^) and HA::MRK1 in MPK2 backgrounds (MPK2^F91F^::myc and MPK2^F91A^::myc). Under basal conditions, the two protein kinases showed limited spatial overlap (Fig 6d, Fig. S6). In particular, MRK1 was not enriched within any of the MPK2-positive polar microdomains, including the posterior cell pole and the cytokinetic bridge during cytokinesis (Fig. 6d).

Given that MPK2 inhibition resulted in redistribution of the protein kinase into a diffuse cytoplasmic pattern, we next asked whether this relocalised MPK2 fraction overlapped with MRK1-containing regions. However, inhibited MPK2^F91A^::myc did not acquire MRK1-like localisation patterns, and MRK1 distribution remained unchanged following inhibition of MPK2 (Fig. 6e). Similarly, MRK1 inhibition did not affect the characteristic location or spatial dynamics of MPK2 (Fig. 6f).

Together, these results demonstrate that although MPK2 and MRK1 inhibition produced closely matching cytokinesis defects, the two protein kinases do not function within a shared linear signalling cascade. Instead, they act through distinct spatially segregated pathways that converge independently to coordinate amastigote cell division.

**Figure 6.**
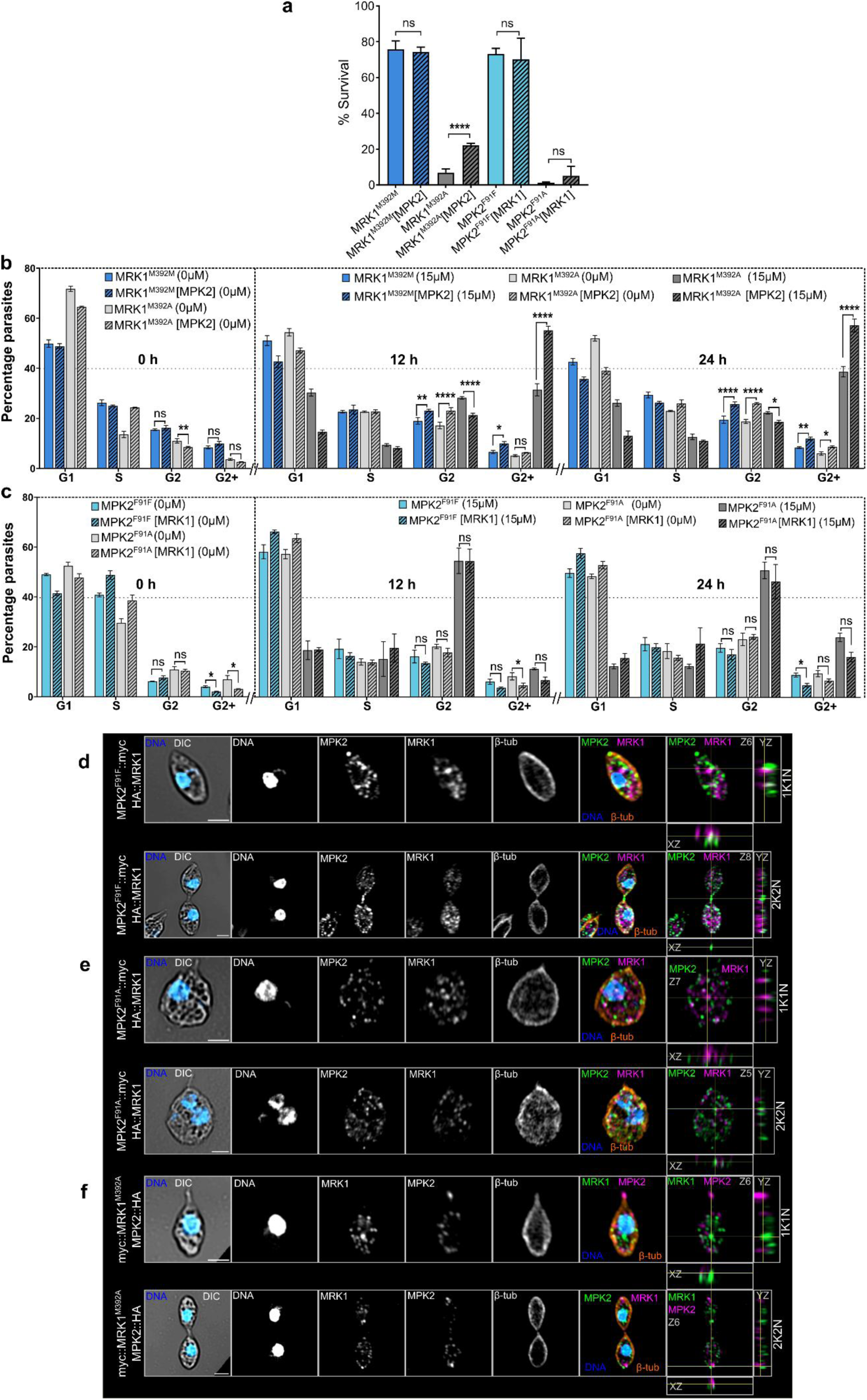
MPK2 and MRK1 operate in separate pathways to regulate cytokinesis. (a) Viability of mature amastigotes of silent and AS mutant lines overexpressing the specified kinases (eg (MRK1^M392M^[MPK2]) and treated with 15 µM 1NA-PP1, assessed by Alamar blue assay after 4 d of replication. Survival was normalised to each cell line’s own DMSO control, set at 100%. Statistical significance was assessed by unpaired two-tailed t-test with Welch’s correction for unequal variances; exact P values are shown for each comparison. (b-c) Cell cycle profiles of silent and AS mutant lines overexpressing the specified kinases with 15 µM 1NA-PP1 or DMSO control, analysed by propidium iodide staining and flow cytometry at 0, 12 and 24 h post-treatment. The percentages of cells in G_1_, S and G_2_ and those with DNA content greater than G_2_ (G_2_+) are reported. Statistical significance was assessed within each cell-cycle phase by two-way ANOVA followed by Sidak’s multiple comparisons test. At 0 h, silent and AS lines were compared to their respective overexpression counterparts (eg MRK1^M392M^ vs MRK1^M392M^[MPK2]; MRK1^M392A^ vs MRK1^M392A^[MPK2]. At 12 h and 24 h, the same comparisons were made separately for untreated and 1NA-PP1-treated pairs. Data represent mean ± SD of three biological replicates; exact P values are provided in Source Data. ns, not significant; *P < 0.05; **P < 0.01; ***P < 0.001; ****P < 0.0001. (d-f). High resolution confocal immunofluorescence micrographs displaying the spatial organisation of HA::MRK1 in (d) MPK2^F91F^::myc, (e) MPK2^F91A^::myc; and of MPK2::HA in (f) myc::MRK1^M392A^; in mature amastigotes after 12 h of treatment with 15 µM of 1NA-PP1. Representative micrographs representing 1K1N (top) and 2K2N (bottom) cells are shown for each background. Merged on the indicated Z-slice and orthogonal views of the full Z-stack (xz and yz cross-sections at the yellow crosshair) are shown. Comparable localisation patterns were observed for HA::MPK2 in the myc::MRK1^M392M^ background (Fig.S6). Protein kinases were labelled with anti-myc (green) and anti-HA (magenta), microtubules were stained with anti-β-tubulin (orange), and DNA was counterstained with DAPI (blue). Scale bar, 2µm.

### Flagellar motility buffers promastigotes against loss of MPK2 activity

The finding that MPK2 and MRK1 activities independently coordinate amastigote cytokinesis raises a fundamental question: why is the requirement for these protein kinases restricted to the amastigote stage, given that cytokinesis is an essential cell cycle event in different replicating life cycle stages of *Leishmania*?

To address this, we first evaluated the requirement for MPK2 and MRK1 activities in promastigotes using the AS kinase mutants. Both MPK2^F91A^ and MRK1^M392A^ displayed increased sensitivity to 1NA-PP1 compared with their corresponding silent controls, with IC_50_ values of 7.49 µM and 3.12 µM respectively, whereas MPK2^F91F^ and MRK1^M392M^ controls showed IC_50_ values of 17.16 µM and 17.96 µM, respectively (Fig. 7a). MPK2^F91A^ showed a 2.3-, 9.5- and 6.3-fold increase in sensitivity to 1NA-PP1 relative to MPK2^F91F^ in promastigote replication, amastigote differentiation and amastigote replication stages, respectively. In contrast, MRK1^M392A^ displayed a 5.8-, 2.8- and 4.2-fold increase in sensitivity relative to MRK1^M392M^ across the same stages. These results revealed a divergence between gene deletion and acute chemical inhibition phenotypes. While Δ*mrk1* is fully viable during promastigote replication, acute inhibition of MRK1^M392A^ triggered severe fitness defects. This suggests that gene deletion may underestimate protein kinase dependency, due to long term genetic adaptation following gene disruption. In contrast, MPK2^F91A^ displayed a genuine stage-specific dependency during amastigote differentiation and replication, but only a marginal fitness defect in replicating promastigotes.

The reduced dependency on MPK2 activity during promastigote replication raised the possibility that promastigote-specific cellular features may buffer the loss of MPK2 activity. A defining morphological distinction between life cycle stages is the presence of a long motile flagellum in the promastigote, which is retracted to a non-motile remnant in amastigotes. We hypothesised that the mechanical forces generated by flagellar motility might help physically drive cell cleavage in promastigotes, thereby mitigating signalling failures caused by loss of MPK2 activity.

To test this, we generated flagellar motility-defective mutants in both the MPK2^F91F^ and MPK2^F91A^ backgrounds by deleting PF16 (MPK2^F91F^::Δ*pf16,* MPK2^F91A^::Δ*pf16*), which generates parasites with paralysed flagella ^31^. Loss of flagellar motility alone did not reduce parasite viability in either the silent control (MPK2^F91F^::Δ*pf16*) or the MPK2 AS background (MPK2^F91A^*::Δpf16*) (Fig. 7b-c). We then treated these lines with a sublethal concentration of 1NA-PP1 (3.75 µM), which resulted in 70% viability of MPK2^F91A^ promastigotes with functional flagella (Fig. 7a, c). In contrast, when MPK2 was chemically inhibited in MPK2^F91A^ parasites carrying paralysed flagella, cell viability dropped nearly 3-fold, down to 25% (Fig. 7d). No obvious change in motility was observed upon MPK2 inhibition alone. This synergistic loss of fitness indicates that flagellar motility is a critical mechanical buffer that dampens the consequences of reduced MPK2 signalling during promastigote division, explaining the heightened vulnerability of non-motile amastigotes to the loss of this protein kinase.

**Figure 7:**
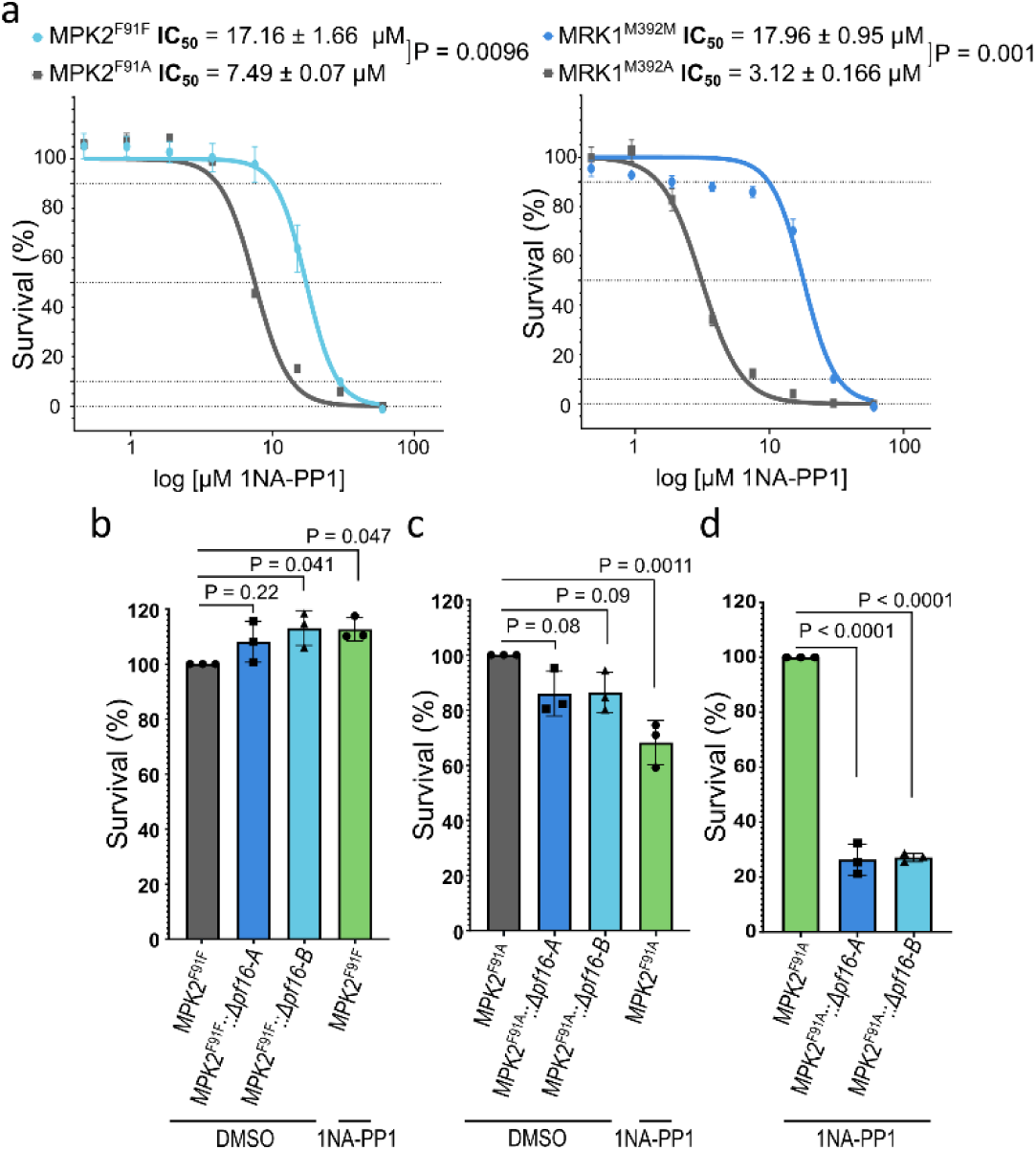
Mechanical buffering of MPK2 activity by the flagellum. (a) Survival of the AS mutants (MPK2^F91A^ and MRK1^M392A^) relative to their respective silent controls (MPK2^F91F^ and MRK1^M392M^), after treatment with 1NA-PP1 during promastigote replication. IC_50_ (mean ± SD) and exact *P* values are shown for each comparison (unpaired two-tailed t-test with Welch’s correction). (b-d) Evaluation of flagellar motility as a buffer for MPK2 inhibition, assessed by Alamar blue viability. Effect of a sublethal dose of 1NA-PP1 (3.75 µM), and flagellar paralysis (two independent *Δpf16* transfection populations -A and -B, untreated) on MPK2^F91F^ (b) and MPK2^F91A^ (c) cell lines. Survival was normalised to untreated MPK2^F91F^ (set at 100%) (b) and MPK2^F91A^ (set at 100%) (c). (d) Combined effect of MPK2 inhibition and flagellar paralysis on MPK2^F91A^ viability. Survival of 1NA-PP1 treated MPK2^F91A^::Δ*pf16* populations was normalised to 1NA-PP1 treated MPK2^F91A^ (with motile flagella), set at 100%. Statistical significance for b, c and d was assessed by one-way ANOVA with Dunnett’s multiple comparisons test. Exact P values are shown for each comparison. Data represent mean ± SD of three biological replicates.

### Time-dependent phosphoproteomic profiling upon MPK2 inhibition

To dissect the signalling cascade downstream of MPK2, we profiled both the phosphoproteome and the proteome of actively replicating axenic amastigotes upon MPK2 inhibition. Analogue-sensitive (MPK2^F91A^) and control (MPK2^F91F^) amastigotes were treated with 15 uM 1NA-PP1 and cells were collected at 0 h (untreated), 4 h and 8 h post-treatment. We identified a total of 7,934 phosphosites and 6,598 proteins in at least three of the four replicates (Table S3). Inhibition triggered specific proteomic shifts, with 71 phosphosites but only 3 proteins differentially regulated 4 and 8 h post-treatment (Table S4, Fig. 8). Notably, two key residues within the MPK2 activation loop, T^174^ and Y^176^, exhibited marked phosphorylation reduction as early as 4 h post-inhibition (log_2_FC = -2.50 and -2.2, respectively), which was sustained through 8 h (Fig. 8a). Sequence alignment confirmed that these residues constitute the conserved TDY dual-phosphorylation activation motif characteristic of MAP kinases (Fig. 8b). The conservation of these phosphorylation sites in *L. donovani* ^20^, suggests that MPK2 activity is required to maintain its own activation in a phosphorylated, fully active state.

Hierarchical clustering of the phosphosites revealed three distinct regulatory patterns (Fig. 8c). Cluster 2 comprised phosphosites that underwent progressive loss of phosphorylation at 4 and 8 h post-treatment and included MPK2 activation loop sites. Functional annotation revealed enrichment in phospoproteins associated with cytoskeletal organisation, flagellum attachment zone (FAZ), microtubule modification and signalling. Notable targets included the cytoskeletal associated proteins CAP156 (S^405^/S^407^S^409^/; log_2_FC = -3.48/-2.31/-2.99) and CAP41, which showed multiple residues lacking phosphorylation (mean log_2_FC = -1.7) after 4 h. Additionally, FAZ14 (S^579^/T^584^; log_2_FC = -1.31/-3.30) and the microtubule depolymerase Kin13-3 (S^460^; log_2_FC = -1.52) also showed phosphorylation reduction. In *T. brucei* homologues of CAP41 and CAP156 localise to the polar cytoskeleton and are important for cleavage furrow placement and cytokinesis ^32,33^. Beyond the cytoskeletal machinery, MPK2 inhibition triggered reduced phosphorylation of signalling hubs, including the essential MAP3K (LmxM.06.0640; S^2490^/S^3091^/S^3095^; Log_2_FC = -2.9/-0.92/-0.41) ^5^ and LST8 (S^296^; Log_2_FC = -2.07 after 8 h), a core component of the TORC1 complex necessary for parasite proliferation and virulence ^23^.

In contrast, Clusters 1 and 3 comprised phosphosites altered predominantly after 8 h of inhibition. Sites in Cluster 1 exhibited an increase in phosphorylation at 8 h in control cells. However, MPK2 inhibition prevented this time-dependent accumulation. Affected sites included the microtubule polyglutamylase TTLL6B (S^1186^, S^1188^, S^1189^; Log_2_FC = -3.6/-3.9/-3.4), the STE kinase LmxM.07.0880 (S^457^, S^467^; Log_2_FC = -2.8/-3.2), the kinetoplastid-specific kinetochore proteins KKT7 (S^83^; Log_2_FC = -2.81) and KKT9 (S^205^, S^206^; Log_2_FC = -2.6/-2) and several RNA binding proteins, including LmxM.17.0990 ( T^1954^/T^1958^; Log_2_FC = -4.7/-5.3). Conversely, Cluster 3 contained sites that lost their phosphorylation at 8 h in control cells, whereas MPK2 inhibition prevented this loss. Proteins in Cluster 3 included the phosphatase NIF (S^244^; Log_2_FC = 4.06), the cytoskeletal proteins CAP51 (Y^96^; Log_2_FC = 1.76) and FAZ12 (S^500^, Log_2_FC = 2.79), TTLL1 (S^554^, S^558^; Log_2_FC = 2.51/2.85), KKT4 (S^509^/S^511^; log_2_FC = 1.35/ 1.73) and the histone lysine crotonylation protein YEATS (T^1126^; log_2_FC = 2.58 at 4h). Overall, regulated proteins in both clusters were predominantly involved in post-transcriptional regulation and nuclear processes.

To confirm the target specificity of these phosphoproteomic responses, we analysed total protein abundance and demonstrated a highly stable proteome, with only three proteins (a cathepsin, a lipase and a hypothetical protein) showing significant changes upon MPK2 inhibition (Table S4).

Overall, the temporal hierarchy of these events demonstrates that under basal conditions, MPK2 maintains activation loop autophosphorylation. Upon MPK2 inhibition, however, a primary loss of phosphorylation of cytoskeletal and FAZ components (Cluster 2) precedes the broader modulation of RNA metabolism and nuclear proteins (Clusters 1 and 3) observed at 8 h.

**Figure 8:**
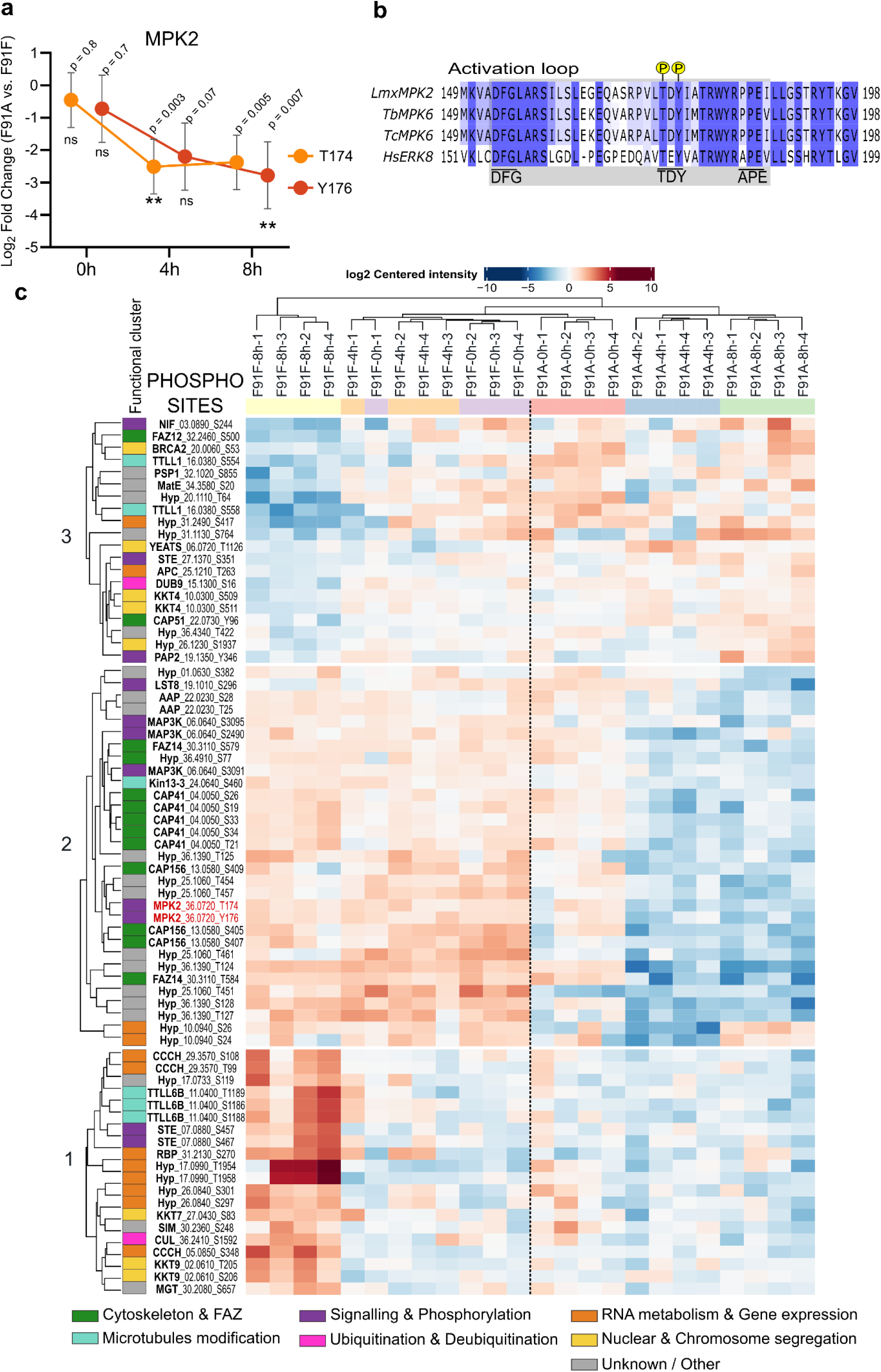
Quantitative phosphoproteomics reveals MPK2 signalling network during axenic amastigote replication. (a) MPK2 phosphorylation kinetics. Changes in phosphorylation levels for Thr^174^ and Tyr^176^ residues in MPK2^F91A^ relative to the MPK2^F91F^ control at 0, 4 and 8 h post-1NA-PP1 inhibition, expressed as log_2_ Fold Change. Points represent the mean and error bars indicate 95% confidence intervals (CI) calculated using DEP2. Significance was defined as |log_2_FC > 1| and adj. P <0.05. (b) Conservation of the TDY motif in the MPK2 activation loop. Multiple sequence alignment of *L. mexicana* MPK2 and its homologues in *T. brucei* (Tb927.10.5140, MPK6), *T. cruzi* (TcCLB.506007.40, MPK6) and *H. sapiens* (Q8TD08, ERK8) performed with MAFFT, default parameters, for visualisation purposes. Conserved catalytic motifs DFG, TDY and APE are highlighted. *L. mexicana* MPK2 phosphorylation sites are indicated by yellow circles above the sequence. (c) Heatmap of differentially modulated phosphosites. Hierarchical clustering of the 71 phosphosites showing significant changes (|log_2_FC > 1| and adj. P <0.05) in MPK2^F91A^ vs. MPK2^F91F^at 4 and 8 h post-1NA-PP1 inhibition (4 biological replicates per condition). Data represent the log_2_ centered intensities, where red indicates increased and blue indicates decreased phosphorylation levels. Functional clustering of the regulated proteins is indicated by the colour-coded bar on the left.

## DISCUSSION

Reversible protein phosphorylation is a major driver of cellular adaptation in kinetoplastid parasites, where polycistronic transcription requires both post-transcriptional and post-translational mechanisms to coordinate morphogenesis and proliferation. Whilst our functional genetic screening of the *L. mexicana* kinome identified stage-essential kinases ^5^, gene deletions cannot distinguish whether a protein kinase is required for both life-cycle stages, specifically during the differentiation process, or for the maintenance of the mature amastigote. By integrating time-course phosphoproteomics across seven differentiation time points with analogue-sensitive chemical genetics, super-resolution microscopy, and substrate profiling, this study provides a systems-level framework linking dynamic phosphorylation cascades to stage-specific cellular architecture and division.

### Dynamic phosphoproteomics maps metabolic and signalling changes during differentiation

Our temporal phosphoproteome expands the catalogue of *Leishmania* phosphorylation dynamics to 16,495 phosphorylation sites (5,795 dynamically regulated), offering phosphosite-specific temporal resolution beyond previous studies ^6,7,9,34^. The timing of these phosphorylation changes is concordant with the metabolic reprogramming during promastigote-to-amastigote conversion. Lower levels of phosphorylation of glucose transporters and enzymes of carbohydrate metabolism within the promastigote clusters, is consistent with the down-regulation of glucose consumption in amastigotes^35^. Conversely, amastigote-associated clusters exhibited elevated phosphorylation of enzymes of phospholipid catabolism and transport, and glutamine biosynthesis, in line with the increased catabolism of fatty acids and the requirements for *de novo* glutamate and glutamine biosynthesis for amastigote growth ^35^. The amastigote-associated clusters were further enriched in acidocalcisome and mitochondrial outer membrane components alongside TORC3 signalling, in line with the reported role of TOR3 in acidocalcisome biogenesis and metabolic regulation ^14^. This suggests that stage-specific metabolic adaptation is hard-wired by protein kinase signalling networks, rather than driven solely by nutrient availability ^35^.

Most developmental phosphosites were mapped outside catalytic domains to intrinsically disordered regions (IDRs), which act as flexible hubs modulating protein-protein interactions, conformational ensembles and protein function ^36^. However, a subset localised to conserved activation loop residues in essential stage-regulated protein kinases ^37^. Phosphorylation of PKAC1 (T^214^) and PKAC3 (T^169^) in the activation loop decreased during differentiation, consistent with reduced activity of these kinases in amastigotes, whereas phosphorylation in AEK1 (S^212^) increased, consistent with enhanced activation. Notably, the corresponding residue in *T. brucei* AEK1 (S^229^) is required for parasite viability ^38^. Similarly, MPK1 (T^176^-Y^178^), MPK2 (T^174^-Y^176^) and MPK15 (T^244^-Y^246^) showed increased phosphorylation of their conserved activation-loop TXY motif, a regulatory element essential for MAPK activation ^3^^9^. Importantly, chemical inhibition of MPK2 induced a rapid loss of phosphorylation of these exact TXY residues, directly linking MPK2 catalytic activity to activation-loop dynamics. The amastigote-specific hyperphosphorylation of the MAPK activation motifs correlates with the loss-of-fitness phenotypes observed in null mutants during differentiation, suggesting that these kinases are catalytically activated during amastigote establishment.

### MPK2 and MRK1 act through independent pathways to coordinate amastigote cytokinesis

MPK2 and MRK1 gene deletions, combined with acute inhibition of the analogue-sensitive MRK1^M392A^ and MPK2^F91A^ lines during differentiation into amastigotes and subsequent replication, displayed strikingly similar phenotypes. These were characterized by the accumulation of multinucleated cells despite undergoing flagellum retraction and cell body rounding. These observations indicate that loss of function of either protein kinase impairs the completion of cell division rather than the differentiation program itself. Cytokinesis in kinetoplastids is a highly coordinated process requiring integration of cell polarity, cytoskeletal organisation and cell-cycle progression ^10,30,40^. In *T. brucei*, disruption of the canonical anterior-to-posterior cytokinesis pathway triggers an alternative, slower, posterior-to-anterior route ^41^. Notably, we observed that while *Leishmania* amastigotes predominantly undergo anterior-to-posterior furrowing, they can naturally execute posterior-to-anterior division. However, inhibition of either MPK2 or MRK1 shifted the preference toward the latter route, accompanied by the dysregulation of nucleus/kinetoplast segregation. MPK2 localises to the anterior and posterior cell poles during division, and its acute inhibition leads to dispersal of the protein kinase into the cytoplasm, indicating that its catalytic activity is required to maintain this polar localisation. Together, these results indicate that MPK2 and MRK1 kinase activities are required to preserve directional fidelity during cell division, highlighting the intrinsic plasticity of the *Leishmania* cytokinesis program.

Despite these convergent phenotypic outcomes, chemical genetics with analogue-sensitive mutants uncovered fundamental functional differences between these two protein kinases. During promastigote proliferation, gene deletions of either MPK2 or MRK1 were well tolerated. However, acute chemical inhibition revealed distinct requirements, as MRK1^M392A^ inhibition caused cell death, whereas MPK2^F91A^ inhibition caused only a marginal fitness defect. This contrast reveals an essential catalytic requirement for MRK1 in promastigotes that was masked in the constitutive gene deletion mutant (Δ*mrk1)*. This suggests that Δ*mrk1* promastigotes likely adapted through compensatory changes to maintain proliferation, a tolerance mechanism that nonetheless proved insufficient during differentiation into amastigotes and subsequent replication. Furthermore, the absence of reciprocal genetic compensation coupled with their distinct subcellular localisations, strongly supports the conclusion that MPK2 and MRK1 act through distinct pathways to execute cytokinesis.

### Substrate profiling links MPK2 to cytoskeletal remodelling

Our temporal MPK2 substrate profiling places MPK2 catalytic activity at the head of a hierarchical cytoskeletal remodelling cascade. Cytoskeletal reorganisation during *Leishmania* cell division involves critical steps, including flagellar pocket duplication and restructuring, symmetrical membrane infolding and subsequent furrow ingression accompanied by daughter-cell twisting and extensive microtubule reconfiguration ^10,42^. Although the molecular mechanisms coordinating these events remain poorly understood, the localisation of MPK2 at *β*-tubulin-positive microtubule tips and within the cytokinetic bridge places the kinase at sites of active cytoskeletal remodelling. Consistent with this, our temporal phosphoproteomic analysis identified multiple cytoskeletal-related proteins as potential MPK2 substrates, as they displayed reduced phosphorylation upon MPK2 inhibition (4 h), including CAP41, CAP156, FAZ14 and KIN13-3. Following 8 h of inhibition, TTLL6B also exhibited a loss of phosphorylation, whereas CAP51, FAZ12 and TTLL1 showed hyperphosphorylation. Among these candidates, CAP41 and CAP51 were previously identified as candidate interacting partners of the KHARON (KH) complex 2 ^43^. The KH complex 2, together with their associated proteins KHAP1 and KHAP2, localises to subpellicular microtubules, where deletion of individual complex components results in multinucleated cells and cytokinesis failure specifically during intracellular amastigote replication ^43^. Other MPK2 regulated proteins provide complementary links to microtubule organisation: FAZ14 and FAZ12 localise to the anterior horseshoe structure at the *Leishmania* flagellar emergence site ^44^, while *L. major* TTLL6B localises at the posterior cell tip. It also possesses microtubule polyglutamylase activity *in vitro* ^45^. Together, these observations suggest that MPK2-dependent phosphorylation contributes to the organisation and remodelling of cytoskeletal structures required for amastigote cytokinesis.

The altered N/K configurations observed following MPK2 or MRK1 inhibition suggests that perturbation of cytokinesis directionality is accompanied by defects in the coordination of organelle segregation. This is consistent with the broader organisation of kinetoplast cell division, in which kinetoplast and nuclear segregation are tightly coordinated with flagellar pocket and cytoskeletal remodelling ^10^. When MPK2 is inhibited, the early dominant response involved a lack of phosphorylation of cytoskeletal proteins, whereas later changes (8 h) affected proteins of RNA metabolism, gene expression and chromosome segregation. Among these were the kinetoplast-specific kinetochore proteins KKT4, KKT7 and KKT9, as well as multiple RNA-binding and nuclear proteins. *L. mexicana* KKT4 possesses microtubule-binding capacity ^46^ and, together with KKT7 and KKT9, exhibits phospho-modulation when cytokinesis is blocked in promastigotes ^47^. Depletion of *T. brucei* KKT4, KKT7 and KKT9 leads to lagging kinetochores during mitosis impairing chromosome segregation fidelity ^46,48^. Given their delayed modulation relative to the early cytoskeletal response, these changes are more consistent with downstream consequences of impaired cell division and cell-cycle progression than with primary MPK2-dependent events. This temporal hierarchy suggests that nuclear and kinetoplast segregation defects are secondary consequences of impaired cytoskeletal remodeling and furrow ingression, rather than primary mitotic failures.

### Flagellar motility acts as a mechanical buffer during parasite division

MPK2^F91A^ displayed a marginal fitness defect during promastigote replication, closely resembling the Δ*mpk2* phenotype, prompting us to investigate why MPK2 activity was required in the amastigote, despite cytokinesis being essential throughout the replicative *Leishmania* life cycle. We hypothesised that mechanical forces generated by flagellar motility might physically facilitate cell cleavage in promastigotes, buffering the parasite against MPK2-dependent signalling impairments. We tested this hypothesis by generating flagellar-motility defective mutants through deletion of the flagellar central pair-associated protein PF16. This deletion generates morphological normal-looking promastigotes with paralysed flagella that remain capable of sustaining infections in mice ^49^, indicating that flagellar beating alone is not essential for parasite replication. Crucially, loss of flagellar motility rendered promastigotes sensitive to MPK2 inhibition, indicating that flagellar motility modulates the cellular requirements for MPK2-dependent cytokinesis. Thus, rather than being an essential driver of cytokinesis, flagellar motility provides functional resilience to loss of MPK2 activity, allowing promastigotes to tolerate impaired MPK2-dependent signalling. In agreement with its essential role in host survival ^5,19^, MPK2 has also been implicated in promastigote adaptation to environmental stress, including responses to arginine deprivation and aquaglyceroporin (AQP1)-dependent antimonite [Sb(Ⅲ)] uptake ^19,20,50^, however, AQP1 phosphorylation was unchanged by MPK2 inhibition in our data. These distinct roles likely reflect the pleiotropic nature for MPK2, acting as a multi-functional signalling node that independently coordinates these cellular processes.

Overall, our work provides a temporal signalling atlas of *Leishmania* differentiation and reveals that protein kinase essentiality is shaped not only by developmental state, but by an interplay between signalling networks and cellular biomechanics.

## METHODS

### Promastigotes cell culture and axenic amastigote differentiation

*L*. *mexicana* (MNYC/BZ/62/M379) promastigotes were grown in HOMEM (Gibco) supplemented with 10% v/v heat-inactivated Fetal Bovine Serum (FBS) (Gibco) and 1% v/v Penicillin/Streptomycin (Sigma-Aldrich). Axenic amastigote differentiation was induced by incubating D3 stationary-phase promastigotes in Schneider’s Drosophila Medium (Sigma-Aldrich) supplemented with 150 μg mL⁻¹ hemin and 20% heat-inactivated FBS, at 33 °C and 5% CO₂. Parasites were subsequently maintained by subculturing cells into fresh medium. Selection drugs were added to the medium as appropriate: 10 μg mL^-1^ blasticidin (InvivoGen), 40 μg mL^-1^ puromycin (InvivoGen), 10–15 μg mL^-1^ G418 (InvivoGen), 50 μg mL^-1^ hygromycin (InvivoGen) and 50 μg mL^-1^ nourseothricin (Jena Bioscience), 15 μg mL^-1^ G418 (InvivoGen).

### *L. mexicana* genetic modifications

Gene deletions and endogenous locus tagging were performed using the CRISPR-Cas9 genome editing toolkit for kinetoplasts ^31^. Primers for sgRNA DNA and donor templates were designed using the LeishGEdit platform (http://leishgedit.net/) and PCR amplified products were electroporated into log-phase promastigotes expressing Cas9 and T7 polymerase (T7Cas9). Drug selection was applied 12 h post-transfection. Clonal lines were established by limiting dilution or maintained as polyclonal populations, as detailed in Table S5. Genomic integration of the tags or gene deletions were verified by diagnostic PCR. To generate complemented lines for null mutants and episomal overexpression lines, the MPK2 or MRK1 coding sequences (CDS) were amplified from *L. mexicana* genomic DNA using primers designed with the NEBuilder Assembly Tool (NEB). For genomic complementation, constructs were assembled into the pRIB-G418 integrative vector (pGL2398), targeting the SSU rRNA locus. For episomal overexpression, the CDS were cloned into pNUS-G418-Ty1 (pGL2944) to obtain Ty1-tagged proteins (N-terminal MRK1; C-terminal MPK2). Assembly reactions were performed using HiFi DNA Assembly Master Mix (NEB), and all constructs were verified by Sanger sequencing. For complementation, pRIB plasmids were linearized with *Pac*I and *Pme*I, and 5 μg of purified DNA was transfected into log-phase promastigotes by electroporation. Selection with G418 was applied after 12h, and clonal lines were established by limiting dilution. Genomic integration in complemented lines were verified by diagnostic PCR. For episomal overexpression, 5 µg of pNUS plasmid was transfected into log-phase promastigotes, maintained as populations under G418 selection, and expression was confirmed by western blot. Precision edits were designed manually and performed as previously described ^51^. The gatekeeper phenylalanine residue at position 91 in MPK2 and the methionine 392 in MRK1 ^26^ were mutated to alanine to generate analogue-sensitive protein kinase mutants (MPK2^F91A^; MRK1^M392A^). As control for genome editing, we generated corresponding silent mutant lines containing identical recoded regions but retaining the original amino acid residues (MPK2^F91F^; MRK1^M392M^). Edits were confirmed by Sanger sequencing (Fig. S4). All transfections were performed using an Amaxa 4D transfection kit and Amaxa 4D nucleofector system (Lonza). A complete list of transgenic cell lines, selection markers, and oligonucleotides used in this study is provided in Table S5.

*L. mexicana* M379 differentiation for phosphoproteomics:

M379 *L. mexicana* promastigotes were cultured in HOMEM (Gibco) supplemented with 10% v/v heat-inactivated FBS (Gibco) and 1% v/v Penicillin/Streptomycin (Sigma-Aldrich) supplemented with 10% v/v heat-inactivated Fetal Bovine Serum (FBS) (Gibco) at an initial density of 1 x 10^5^ parasites mL^-1^. After 2 d, parasites reached logarithmic phase (5 x 10^6^ parasites mL^-1^) at which point the first samples were collected (LOG). The subsequent day, marking the cessation of replication, was considered the first day of stationary-phase. The culture was maintained in stationary-phase for 3 d before sampling (STAT). To induce differentiation, 3 d stationary phase promastigotes were harvested, washed and resuspended in Schneider’s Drosophila medium supplemented with 20% FBS (Gibco) 1% Penicillin/Streptomycin and 150 μg mL⁻¹ , adjusted to pH 5.5 and incubated at 33 °C. Samples were collected at 2h (AxA2), 8h (AxA8), 24h (AxA24), 48h (AxA48) and 5 days (AxA5D) post-induction. For each time point, 10 biological replicates were collected and to ensure a comprehensive representation of the phosphoproteome, a dual-lysis strategy was employed: five replicates were lysed in Urea lysis buffer (20 mM HEPES pH 8, 9 M urea) to facilitate the recovery of soluble proteins, while the remaining five were lysed in SDS lysis buffer (5% SDS, 50 mM TEAB pH 8.5) to enhance the solubilisation of cytoskeletal proteins. Both buffers were supplemented with protease and phosphatase inhibitors (1 mM sodium orthovanadate, 2.5 mM sodium pyrophosphate, 1 mM β-glycerophosphate, 1x HALT, 1x Complete Roche protease inhibitor EDTA free, 1 mM E64d and 1x PhosSTOP).

### MPK2 inhibition for phosphoproteomics

MPK2^F91F^ and MPK2^F91A^ cell lines were differentiated to axenic amastigotes. On d 4, cells were treated with 15 µM 1NA-PP1 (4-Amino-1-tert-butyl-3-(1’-naphthyl)pyrazolo[3,4-d]pyrimidine; Toronto Research Chemical) or DMSO control and samples were collected at 0 h (0 µM 1NA-PP1), 4 h (15 µM 1NA-PP1) and 8 h (15 µM 1NA-PP1) after treatment. For each time point, 5 biological replicates were collected and lysed in SDS lysis buffer (5% SDS, 50 mM TEAB pH 8.5) to enhance the solubilisation of cytoskeletal proteins, supplemented with protease and phosphatase inhibitors (1 mM sodium orthovanadate, 2.5 mM sodium pyrophosphate, 1 mM β-glycerophosphate, 1x HALT, 1x Complete Roche protease inhibitor EDTA free, 1 mM E64d and 1x PhosSTOP).

### Phosphoproteomics of *L. mexicana*

For the M379 differentiation and the MPK2 inhibition analyses, samples were digested in parallel from SDS-and urea-lysed cells, using S-Trap and in-solution digestions, respectively. Protein was first reduced with 5.7 mM tris(2-carboxyethyl)phosphine and heated to 55 °C for 15 min before alkylation with 22.7 mM methyl methanethiosulfonate at room temperature for 10 min. For S-Trap digestion of SDS-lysed samples, protein was acidified with 6.5 mL of aqueous 27.5 % (v:v) phosphoric acid then precipitated with dilution seven-fold into 100 mM TEAB 90% (v:v) methanol. Precipitated protein was captured on S-trap (Profiti – C0-micro) and washed five times with 165 µL 100 mM TEAB 90% (v:v) methanol before digesting with the addition of 20 µL of 0.1 µg µL^-1^ Promega Trypsin/Lys-C mix (V5071) in aqueous 50 mM TEAB and incubation at 47 °C on hot plate for 2h. Peptides were recovered from S-trap by spinning at 4000 *g* for 60 s. S-traps were washed with 40 µL aqueous 0.2% (v:v) formic acid and 40 µL 50% (v:v) acetonitrile:water and washes combined with the first peptide elution. For in-solution digestion of urea-lysed samples, protein solutions were diluted to 2 M urea with aqueous 50 mM ammonium bicarbonate, before adding Promega Trypsin/Lys-C mix (V5071) at 1:50 protease to protein ratio and incubation at 37°C for 16 h. Peptides were desalted using Pierce graphite spin columns (Thermo Scientific – 88302) following the manufacturer’s procedure. From each digested pool of peptides 95% of the material was taken for phosphopeptide enrichment and 5% held for total protein analysis. Phosphopeptides were enriched using titanium dioxide-functionalized magnetic microspheres (MagReSyn - MR-TID02) following the manufacturer’s protocol. Briefly, peptides were resuspended in 1 M glycolic acid in 80% (v:v) acetonitrile (ACN) and 5% (v:v) trifluoroacetic acid (TFA) and mixed with 200 mL of equilibrated microspheres. Bound phosphopeptides were washed with aqueous 80% (v:v) ACN containing 1% (v:v) TFA, then aqueous 10% (v:v) ACN containing 0.2% (v:v) TFA, before eluting with aqueous 1% (v:v) ammonium hydroxide. Phosphopeptide-enriched eluent was dried down in a vacuum concentrator and reconstituted in aqueous 0.1% (v:v) TFA for LC-MS/MS.

Peptides were loaded onto EvoTip Pure tips for desalting and as a disposable trap column for nanoUPLC using an EvoSep One system. Pre-set EvoSep gradients were used with a 15 cm EvoSep C_18_ Performance column (15 cm x 150 mm x 1.5 mm). Non-enriched peptides for total proteomics were separated over a 30 SPD gradient and enriched phosphopeptides were eluted over a 15 SPD gradient. The nanoUPLC system was interfaced to a timsTOF HT mass spectrometer (Bruker) with a CaptiveSpray ionisation source (Source). Instrument source settings were: capillary voltage, 1,700 V; dry gas, 3L min^-1^; dry temperature; 180°C. NanoESI-MS and MS^2^ spectra were acquired using Compass HyStar software (version 6.2, Bruker). MS spectra were acquired between *m/z* 100-1,700. TIMS settings were: 1/K0 0.6-1.60 V.s/cm^2^; Ramp time, 100 ms; Ramp rate 9.42 Hz. Non-enriched peptides for total proteomics were acquired in positive PASEF-DIA mode, with DIA windows of 26 Th width between *m/z* 400-1201 and a TIMS range of 1/K0 0.6-1.60 V.s/cm^2^. Phosphopeptides were acquired in positive PASEF-DDA mode, with 10 PASEF ramps per cycle and a total cycle time of 1.9 s. An intensity threshold of 2,500 and a target intensity of 20,000 were set with active exclusion applied for 0.4 min post precursor selection. For both DIA and DDA acquisitions, collision energy was interpolated between 20 eV at 0.65 V.s/cm^2^ to 59 eV at 1.6 V.s/cm^2^.

DIA spectra were searched using Spectronaut (Biognosys – v20.4). DDA data were searched using FragPipe (v23.1) ^52^. Searches were performed against the LmexCas9T7-prot_2021.fasta ^53^ database appended with common proteomic contaminants. DIA search criteria specified: enzymes, trypsin/P; variable modifications; acetyl (protein N-term), oxidation (M); fixed modification, methylthio (C); digest type, specific; max peptide length, 52; min peptide length, 7; missed cleavages, 2; MZ extraction strategy, maximum intensity; precursor Q Value cutoff, 0.01. DIA XIC extraction windows for IM, IT, MS^1^ and MS^2^ were all set as dynamic. Identifications were filtered to require protein q-values < 0.01 and a minimum of two peptides per accepted protein. DDA search criteria specified trypsin/P protease with up to two missed cleavage positions. Mass tolerance was set to 15 ppm for MS^1^ and MS^2^. For variable modifications, oxidation of methionine, protein N-terminal acetylation and phosphorylation (S,T,Y) were selected. Validation was performed using ProteinProphet and PeptideProphet within Philosopher (v5.1). Searches were run at 1% FDR for both peptide and protein as assessed empirically against a reverse database search. Site localisation probabilities were calculated with PeptideProphet. Intensities from accepted peptides were extracted using IonQuant (v1.11).

### Data processing, differential abundance analysis and cluster analysis

Label-free intensities were exported from Spectronaut (7,051 protein groups identified) and FragPipe (44,561 phosphosites identified) and processed using DEP2 and limma R packages. A total of 7,051 protein groups and 16,495 phosphosites (in 3,101 proteins) identified in 3 out of 5 biological replicates in at least one experimental condition were retained. Raw intensities were normalised using the Variance Stabilizing Transformation (VSN) method. Missing values were imputed using the Minimum Probability (MinProb, q = 0.01) method as missing values were assumed to be predominantly missing not at random (MNAR). Sample reproducibility was assessed by principal component analysis and sample-to-sample correlation (Fig. S1). For the differential abundance analysis, we defined LOG as control while multiple testing correction was performed using the Benjamini-Hochberg (BH) False Discovery Rate (FDR) procedure. Features were considered significantly regulated if they exhibited an adj. *P* < 0.05 and a |log_2_ FC| > 1. Under these criteria, 5,795 phosphosites (in 1,888 proteins). The fold-change cut-off excluded no additional phosphosites. Imputed values represented 57.4% of the filtered phosphoproteome and 3.1% of the total proteome. For the phosphoproteome, two checks were performed: the significance rate was lowest among sites quantified in a single condition (16.6%) and highest among those quantified in six conditions (67.9%); and in 96.9% of the conditions in which a significant phosphosite was not detected, its parent protein remained quantified in the total proteome. Complete results are provided in Table S1.

Significant phosphosite log2 fold-change profiles (relative to the LOG time point) were standardised (*standardise*) and clustered using the fuzzy c-means algorithm (Mfuzz; seed 123). The fuzzifier was estimated by *mestimate* (*m* = 1.55). The number of clusters was set to 10, the largest value at which no two clusters centroids were correlated at r > 0.9; redundant profiles appeared from c = 11 onwards. At k = 10, 64.2% of phosphosites had a maximum membership score (α) ≥ 0.5. Each of the 5,795 significant phosphosites was assigned to the cluster for which it had the highest membership score; membership scores are reported as a measure of assignment confidence and were not used to filter sites. Clusters were renumbered according to the developmental order of their centroids, and this numbering is used throughout.

### GO enrichment

Gene ontology (GO) enrichment (computed and curated) was performed using the gene identifiers from each cluster with the TriTrypDB enrichment tool (release 69), using the complete *L. mexicana* genome as background. Terms with a nominal p-value < 0.05 and a fold enrichment > 1.5 were retained. Broad/generic parent terms, obsolete categories, and semantically redundant terms were systematically excluded. When multiple terms shared the same biological meaning, only the most descriptive and representative category was selected. Additionally, terms with a background count of ≤ 2, a target results count ≤ 1, or those directly inherent to the nature of the dataset (e. g. phosphorylation) were discarded. Curated GO enrichment results are provided in Table S2.

### Total proteome analysis

The total proteome was processed identically to the phosphoproteome (same filter, vsn, normalisation, MinProb imputation with q = 0.01, contrasts against LOG and significance criteria). Protein groups listing several genes were expanded so that each gene inherits the group quantification and test result (7,051 protein groups, 7,294 genes). Protein log2-fold change profiles (relative to LOG) were standardised with the same Mfuzz procedure; ensuring identical scale and variance across both datasets. Rather than clustering the total protein data independently, protein profiles were mapped to their corresponding phosphosites, preserving the cluster structure of the phosphoproteome. Phosphosites whose parent protein was not quantified in the total proteome (n = 43) were excluded. Pearson correlation coefficients (r) between each phosphosite and its parent protein were computed on the standardised seven-time points profiles, and the mean r across phosphosites was reported for each cluster, together with the percentage of parent proteins significantly modulated in abundance.

### Data processing and differential abundance analysis for MPK2 inhibition

Label-free intensities were exported from Spectronaut (6,599 total proteins identified) and FragPipe (17,504 phosphosites identified) and processed using DEP2 and limma packages. Five biological replicates were acquired per condition; replicate 4 was excluded from both datasets owing to low phosphopeptide identification (median 86.2% missing values across conditions, vs. 60.4 - 65.5% for the other replicates). One untreated AS sample (F91A_0h_1) also showed reduced identification but was retained, as this was restricted to a single sample and after imputation it clustered with the other control samples (Fig. S7). Phosphosites (7,934) and protein groups (6,598) quantified in at least 3 of 4 replicates in at least one condition were retained. Data was normalised by variance-stabilizing transformation (vsn). Missing values were imputed with a mixed strategy, features missing in all replicates of at least one condition were considered MNAR and imputed with MinProb (q = 0.01) while the remaining missing values were imputed by k-nearest neighbors (k = 10). Because each contrast compared two different lines, which may differ in protein abundance, phosphosite intensities were normalised to the abundance of their parent protein. For each identified phosphosite, the log_2_ intensity of the “parent” protein was subtracted from the phosphosite value (log_2_ Phospho - log_2_ Total Protein). Out of 7,934 initial phosphosites, 7,860 (99.1 %) were successfully matched to their parent protein and corrected for abundance. The remaining 74 sites, for which the parent protein was not detected in the total proteome dataset, were retained unnormalised. For the differential abundance analysis, we defined manual contrasts to evaluate the effect of MPK2 inhibition at different time points, as detailed in the Results section. Multiple testing correction was performed using the Benjamini-Hochberg (BH) False Discovery Rate (FDR) procedure. Sites and proteins were considered significantly regulated if they exhibited an adj. P < 0.05 and a |log_2_ FC| > 1. Under this criteria, 72 phosphosites (in 44 proteins) and 3 total proteins were significantly modified. Heatmap was generated using the plot_heatmap function in DEP2. Phosphosites significantly regulated in F91A_4h vs. F91F_4h and F91A_8h vs. F91F_8h (71 sites), were displayed as row-centered log_2_ intensities, clustered by k-means (k = 3, selected by elbow criteria), with samples ordered by hierarchical clustering (Euclidean distance). Raw phosphosites and protein intensities are provided in Table S3, whereas regulated sites and proteins are described in Table S4.

### Functional Annotation of data from MPK2 Inhibition

Homologs for *Leishmania mexicana* proteins were identified using the OrthoMCL (version 7) and Synteny tools available in TriTrypDB. Gene Ontology (GO) terms were retrieved from TriTrypDB, incorporating both curated and computed evidence. To expand the functional landscape, enrichment analysis was also performed using DAVID (Database for Annotation, Visualization and Integrated Discovery), mapping *L. mexicana* gene IDs to their closest well-characterized model organisms. To complement the gene ontology analysis, a literature curation was conducted for each protein or its *T. brucei* ortholog, prioritizing experimental evidence related to subcellular localisation and cellular functions. By integrating this information, proteins were manually assigned to one of six Functional Clusters: (1) Cytoskeleton & FAZ, (2) Microtubules modification, (3) RNA Metabolism & Gene expression, (4) Signalling & Phosphorylation, (5) Nuclear & Chromosome segregation and (6) Ubiquitination & Deubiquitination. Proteins with less represented or no clear functional evidence were categorized as “Unknown / Other”.

### 1NA-PP1 susceptibility assays

Dose-response assays were performed in 96-well plates. Parasites were seeded into serial dilutions of the bumped kinase inhibitor 1NA-PP1, ranging from 60 µM to an untreated/DMSO control only. Susceptibility of gatekeeper mutants and their respective silent controls were assessed by addition of 40 µl of 0.0125% (w/v) resazurin (Alamar Blue) prepared in PBS, added 10 h before the end of the treatment period; fluorescence was measured using a CLARIOstar plate reader (BMG LABTECH), excitation 540 nm / emission 590 nm. Survival at each concentration was normalised to each line’s own untreated/DMSO control, set at 100%. Dose-response curves were fitted and IC_50_ values calculated by nonlinear regression (log[inhibitor] vs. normalised response, variable slope) using GraphPad Prism v10.4.1. Three parasite stages were assessed. Firstly, logarithmic-phase promastigotes were seeded at 1 x 10^4^ parasites mL^-1^ and susceptibility was measured after 96 h of treatment. Secondly, stationary-phase promastigotes (day 3 of stationary-phase) were seeded at 1 x 10^5^ parasites mL^-1^ concomitant with induction of differentiation and susceptibility was measured after 120 h of combined differentiation and inhibitor treatment, the same time point used to characterise the null mutant kinases. Thirdly, amastigotes differentiated for 4 days were seeded at 1 x 10^5^ parasites mL^-1^ and susceptibility was measured after 96 h of inhibitor treatment. IC_50_ values for each analogue-sensitive line were compared to its respective silent control by unpaired two-tailed t-test with Welch’s correction for unequal variances, applied uniformly across all comparisons.

### Cell viability assays

Protein kinase null mutants and complemented lines were grown to stationary-phase in complete HOMEM media, and on the third day they were resuspended at 1 x 10 ^5^ cells mL^-1^ in complete Schneider media pH 5.5 and incubated at 33°C, 5% CO_2_ for five days. Analogue-sensitive and silent mutants cross-complemented with the reciprocal kinase differentiated into axenic amastigotes after 4 days in complete Schneider’s *Drosophila* medium pH 5.5, 33°C, and 5% CO_2_ were resuspended at 1 x 10 ^5^ cells mL^-1^ with 15 µM 1NA-PP1 or with the equivalent volume of DMSO (Control), and incubated for another further 4 days in the same media and conditions. MPK2 analogue-sensitive and silent mutants carrying a knock-out of a flagellar motility gene (PF16) were grown to log-phase and resuspended at 1 x 10 ^4^ cells mL^-1^ in complete HOMEM with 3.75 µM 1NA-PP1 or with the equivalent volume of DMSO (DMSO control), and incubated at 26°C for 4 days. Susceptibility was assessed by addition of 40 µl of 0.0125% (w/v) resazurin (Alamar Blue) prepared in PBS, added 10 h before the end of the treatment period. Fluorescence at 590 nm was recorded using the CLARIOstar Plate Reader (BMG Labtech). Data were normalised to the specified control cell line in the different assays and presented as an average of three biological replicates, each with six technical replicates, unless otherwise indicated.

### Cell cycle assessment with flow cytometry

T7Cas9, Δ*mpk2* and Δ*mrk1* cells on the third day of stationary phase were differentiated to axenic amastigotes and samples were collected at stationary phase, 24 and 72 h after induction of differentiation. Analogue-sensitive lines and their respective silent controls were differentiated to day 4 axenic amastigotes, treated with 15 μM 1NA-PP1 for 12 or 24 h, and harvested. For all conditions, cells were washed with PBS containing 5 mM EDTA (PBS-EDTA), fixed in 70% methanol in PBS-EDTA and stored at 4°C until analysis. Cells were then washed with PBS-EDTA and resuspended in 0.5 mL PBS-EDTA containing 10 µg mL^-1^ of propidium iodide (PI) and 10 µg mL^-1^ of RNase A. Following overnight incubation at 4°C, cells were transferred to a 96-well plate and DNA content was measured by flow cytometry using a CytoFLEX LX cytometer (Bechman Coulter). For each cell line, 10,000 events were acquired across 3 biological replicates. Intact parasites were gated using forward scatter height (FSC-H) vs. side scatter height (SSC-H). Doublets were excluded using sequential gating on PI-height vs. PI-area (Y610-mCherry) and FSC-H vs. FSC-A. Cell cycle phase distribution was quantified using the Watson (Pragmatic) model in FlowJo software v10.10.0. The percentages of cells in G_1_, S and G_2_ and those with DNA content greater than G_2_ (G_2_+) automatically calculated by the model were reported. The resulting percentages were plotted and statistical analyses were performed as indicated in the respective figure legend using GraphPad Prism v10.4.1.

### Fixed cells microscopy

*L. mexicana* M379 cells were collected at LOG, STAT, AxA2, AxA8, AxA24, AxA48 and AxA5D timepoints, washed in PBS, fixed in 4 % paraformaldehyde for 10 min at room temperature, washed twice in PBS (1,000 x *g*, 10 min) and incubated with 10 μg mL^-1^ Hoechst 33342 at room temperature for 10 min. Excess Hoechst was removed by washing the cells in PBS and the final pellet was resuspended at approximately 5 x 10^6^ parasites per mL. Suspension was added to a slide and imaged in a Zeiss AxioObserver 7 Inverted Microscope.

### Morphometric analysis

T7Cas9, Δ*mpk2* and Δ*mrk1* cells were imaged as described above 3 days after induction of differentiation. For each line, cells were randomly selected from two independent biological replicates and cell body length and width were measured using ImageJ v1.54r (Fiji) ^54^. Differences relative to the T7Cas9 line were assessed by ordinary one-way ANOVA with Dunnett’s multiple comparison test using GraphPad Prism v10.4.1.

### Live cell imaging

Localisation of mNeonGreen-tagged MPK2 and MRK1 was examined in live stationary-phase promastigotes (STAT) and axenic amastigotes 24 h after differentiation (AxA24). 1 x 10^6^ cells were harvested by centrifugation (1,200 x *g*, 10 min), washed in PBS and incubated with 10 μg mL^-1^ Hoechst 33342 at room temperature for 10 min. Cells were washed in ice-cold PBS and resuspended in 50 μL CyGel (BioStatus) on ice. 25 μL of the suspension was dispensed onto a slide, covered with a coverslip and imaged immediately in a Zeiss AxioObserver 7 Inverted Microscope. Hoechst was imaged with a 385 nm LED and 430-470 nm emission filter, and mNeonGreen with a 475 nm LED and 500-550 nm emission filter. Z-stacks were deconvolved using the Microvolution plugin for Fiji (100 iterations).

### Immunofluorescence microscopy

Cells were differentiated to axenic amastigotes, and after 4 d were treated with 15 μM 1NA-PP1 for 12 h where indicated. Parasites were harvested, washed twice with PBS (1,200 x *g*, 10 min) and allowed to adhere to poly-L-lysine coated coverslips for 20 min at 33 °C. Excess liquid was removed and cells were fixed at room temperature with 4% paraformaldehyde in PBS for 10 min. Following two washes with PBS, cells were permeabilized with ice-cold 100% methanol for 10 min, washed with PBS and blocked with 5% FBS in PBS containing 0.1% saponin for 2 h. Sequential immunostaining was performed using antibodies diluted in blocking buffer, with three PBS washes applied between incubations. The following primary and secondary antibody combinations were used: Rabbit anti-Myc (clone 71D10; Cell signalling Technology; 1:200 dilution, 2h incubation) paired with goat anti-rabbit Alexa Fluor 488 (cat. A-11034; Thermo Fisher; 1:200 dilution, 40 min incubation); Rat anti-HA (clone 3F10; Roche; 1:100 dilution, 2h incubation) paired with donkey anti-rat Alexa Fluor 647 (cat. A78947; Thermo Fisher; 1:250 dilution, 40 min incubation); Mouse anti-β-tubulin (clone KMX-1; Sigma-Aldrich; 1:800 dilution, 1h incubation) paired with goat anti-mouse Janelia Fluor 549 (cat. NBP1-72739JF549; Novus Biologicals; 1:100 dilution, 40 min incubation). Following 3 washes with PBS, cells were counterstained with 20 μg mL^-1^ DAPI in PBS for 30 min, followed by a final PBS wash and coverslips were mounted on glass slides using ProLong glass antifade mountant (Invitrogen). Images were captured in a Zeiss LSM980 MP AiryScan 2 confocal microscope with z-stack slices intervals of 100-150 μm (DAPI Excitation 405 nm Emission 416-484 nm; AF488 Excitation 488 nm Emission 491-553 nm; AF647 Excitation 639 nm Emission 656-755 nm; AF549 Excitation 594 nm Emission 597-641 nm) and post-processed using LSM Plus deconvolution module in Zeiss Zen software.

### Amastigote polarity, cytokinesis and DNA segregation analysis

Parasite polarity, cell cycle status, DNA segregation and protein kinase localisation were quantitatively evaluated across full confocal Z-stacks using combined DIC optics and fluorescence microscopy (DAPI, anti-β-tubulin, anti-myc immunolabeling). The anterior-posterior axis was defined by the position of the kinetoplast relative to the nucleus and the concave/convex curvature of the kinetoplast disc. To assess cell division status, amastigotes without a discernible cleavage furrow and possessing a single nucleus and kinetoplast (1K1N) were classified as *non-dividing*. Parasites were classified as *dividing* if they exhibited duplicated organelles (2K2N, 2K1N, 1N2K) or presented a visible cleavage furrow in both tubulin and DIC channels regardless of organelle count (e.g. enlarged 1K1N cells undergoing cleavage). For dividing cells, the spatial progression of cytokinesis was determined by mapping the site of cleavage furrow initiation relative to the established anterior-posterior axis, categorising it as anterior, posterior or bidirectional cytokinesis. DNA distribution and organelle segregation were subsequently evaluated in dividing parasites. Cells were scored as *normal* when displaying two fully segregated nuclei and two kinetoplasts (2K2N) correctly positioned along the cellular axes. Conversely, cells were categorised as *abnormal* if they exhibited at least one of the following defects: incomplete nuclear or kinetoplast segregation, aberrant copy number (such as 1N2K, 2N1K, multinucleate, or multi kinetoplast), or mislocalisation of nuclear or kinetoplast DNA within the cell body. Subcellular localisation of target protein kinases was scored relative to the defined cell poles (anterior/posterior enrichment) or categorised as cytoplasmic. Statistical significance was assessed using Fisher’s Exact Test on raw counts. Sample sizes (*n* = 51 to 238 per condition) and exact P values are detailed in the Source Data file.

### Western blotting

Parasite pellets were lysed in lysis buffer (8 M Urea, 2% SDS, 50 mM Tris-HCl, pH 7.6) supplemented with protease inhibitors (1x HALT, 1x Complete Roche protease inhibitor EDTA free, 1 mM E64d) and 1 μl of Basemuncher endonuclease (Abcam). Lysates were incubated on ice for 15 min. Protein concentrations were determined using the Pierce BCA Protein Assay Kit (Thermo Fisher) following manufacturer’s instructions, and 100 μg of total protein per sample was prepared for loading. Samples were diluted in NuPAGE LDS sample buffer containing 100 mM dithiothreitol (DTT) and denatured at 75 °C for 10 min. Equal amounts of protein were loaded onto 4-12% NuPAGE Bis-Tris Mini Protein Gels, separated in NuPAGE MOPS SDS Running Buffer, and transferred onto PVDF membranes using the iBlot 2 Gel Transfer System (Invitrogen). Membranes were blocked with 5% skimmed milk in TBS-T for 1 h at room temperature. Membranes were then incubated overnight at 4°C in blocking buffer with primary antibodies: mouse anti-Myc (1:1000; clone 9B11; Cell Signalling) or mouse anti-Ty1 (1:500). Following washes with TBS-T, membranes were incubated with HRP-conjugated goat anti-mouse IgG secondary antibody (1: 5,000 dilution; cat. W4021; Promega) for 1 h at RT in blocking buffer. Protein bands were visualised using Clarity Max Western ECL Substrates (Bio-Rad) and chemiluminescence images were acquired using a Chemidoc MP Imaging System (Bio-Rad). To detect the loading control, membranes were stripped during 15 min using Restore Western Blot Stripping Buffer (Thermo Fisher), re-blocked and incubated overnight at 4 °C with sheep anti-OPB antibody (1:10,000) as a loading control. Membranes were washed with TBS-T, incubated with HRP-conjugated anti-sheep IgG secondary antibody (1:5,000; GeneTex) for 1 h at RT, and developed as described above.

## Acknowledgements

This work was supported by funding from the Wellcome Trust (223045/Z/21/Z). The funders had no role in study design, data collection, data analysis, interpretation, and writing of the manuscript. We thank Rachel Neish for assistance with proteomics sample collection, Mathieu Cayla for sharing initial code templates and Martin Wiese for sharing unpublished phosphoproteomics data. We also thank our expert colleagues in the Bioscience Technology Facility of University of York who assisted our microscopy, flow cytometry, genomic data analysis, and mass spectrometry research. The York Centre of Excellence in Mass Spectrometry was created thanks to a major capital investment through Science City York, supported by Yorkshire Forward with funds from the Northern Way Initiative, and subsequent support from EPSRC (EP/K039660/1; EP/M028127/1).

## Author contributions

**Y. Romina Nievas**: Methodology, Investigation, Visualization, Conceptualization, Writing – original draft, Writing – review & editing. **Charlotte McNiven:** Investigation. **Adam Dowle:** Methodology, Investigation, Data Curation. **Jeremy Mottram:** Writing – review & editing, Supervision, Methodology, Conceptualization, Funding Acquisition.

## Supplementary figures

**Figure S1.**
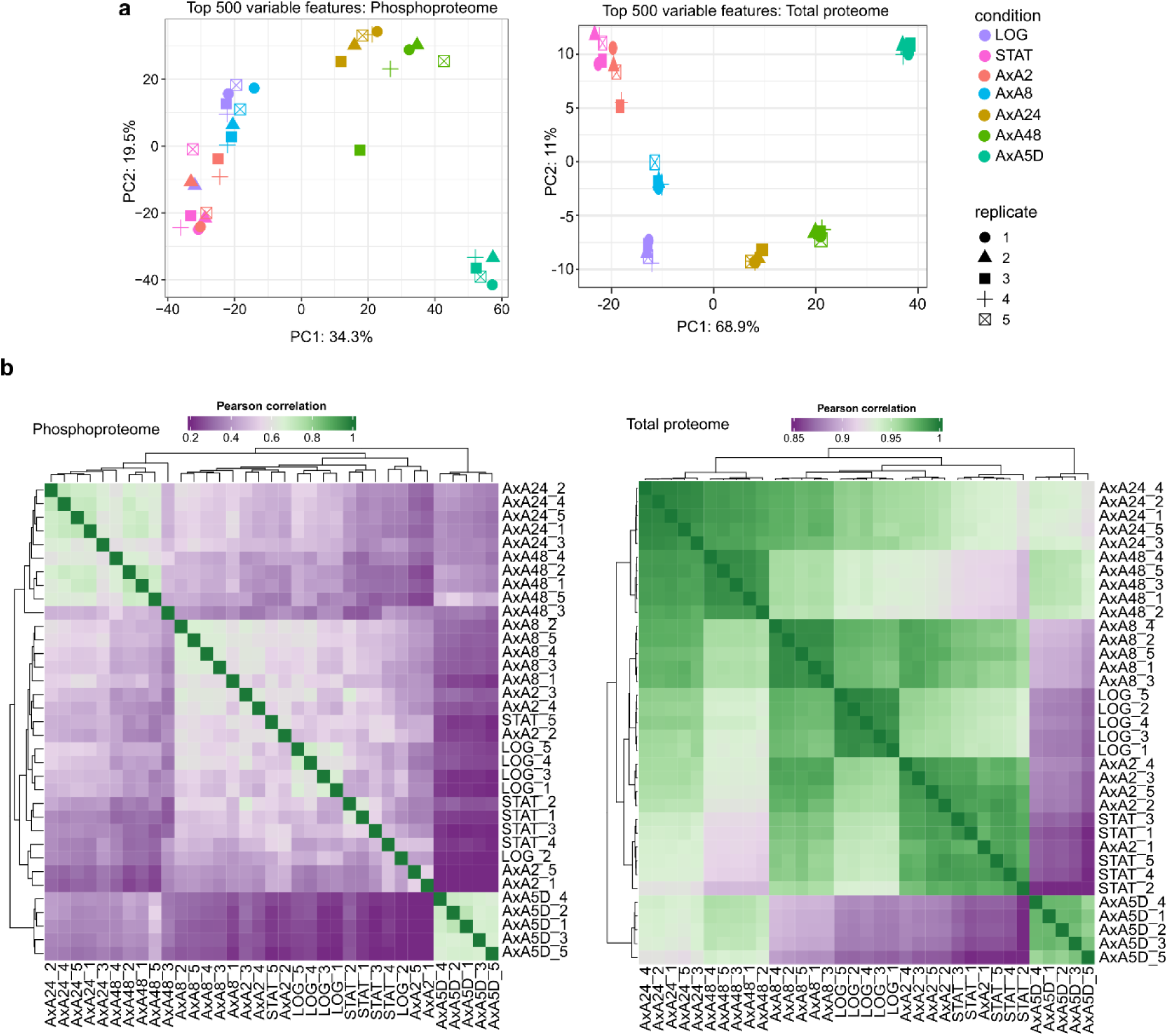
Quality control of differentiation phosphoproteomic and total proteomic dataset. (a) Principal component analysis of the 35 samples, based on the 500 most variable phosphosites or proteins after filtering, normalisation and imputation. Color denotes condition and symbol denotes replicates. (b) Pearson correlation between all pairs of samples for the phosphoproteome (left) and the total proteome (right), with hierarchical clustering of rows and columns.

**Figure S2.**
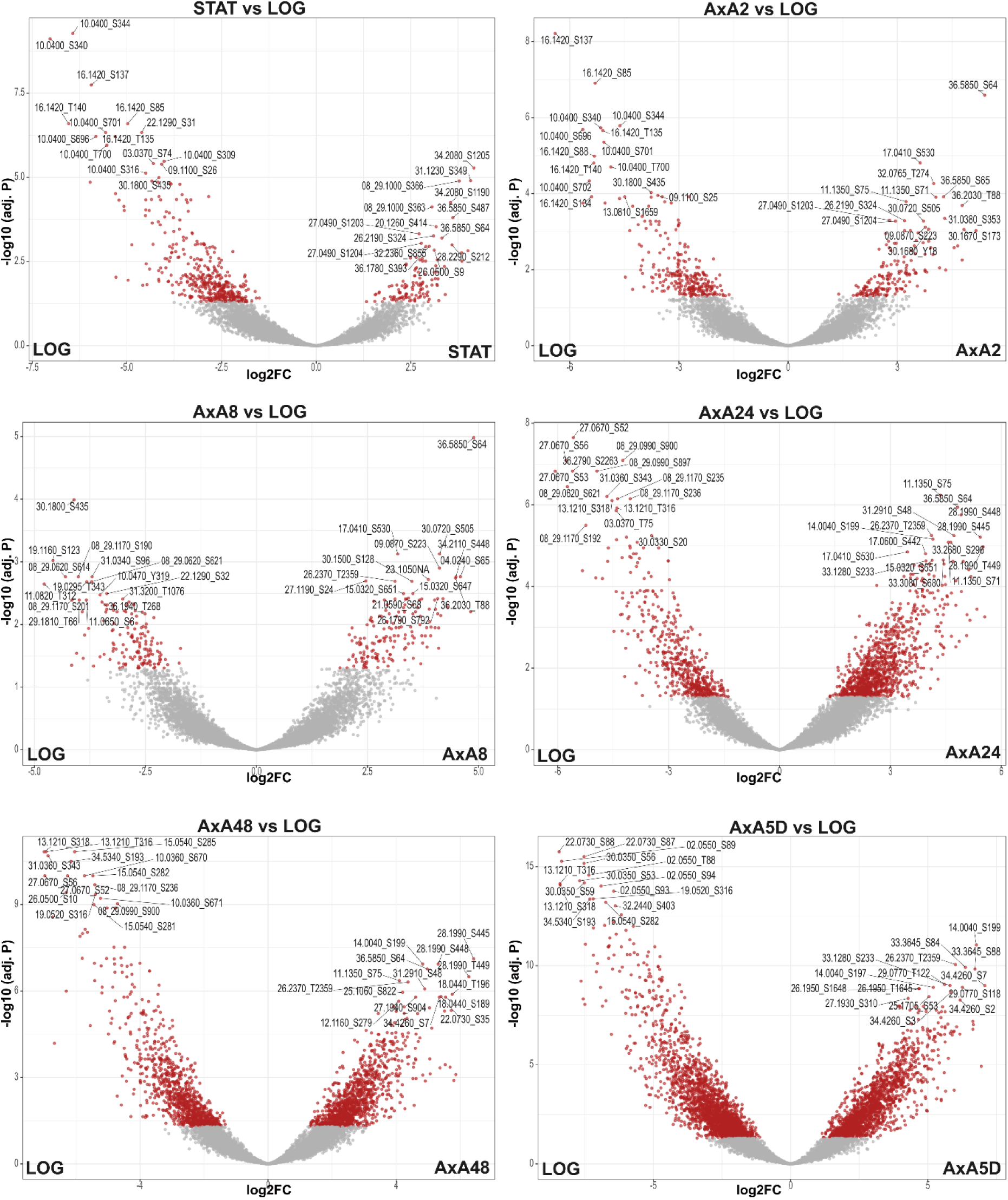
Distribution of modulated phosphorylation sites during differentiation. Volcano plots display phosphosite abundance variations across the differentiation time course, with logarithmic promastigotes (LOG) serving as the reference control. Individual panels represent sequential contrasts (STAT through AxA5D vs. LOG). Sites achieving statistical significance (adj. *P* < 0.05, |log_2_FC| > 1) are highlighted in red, while non-regulated sites are depicted in grey. Within each stage comparison, the 15 most prominent up-regulated and down-regulated phosphosites (prioritised by adjusted *P*-value) are annotated with their respective gene IDs and specific phosphorylated residues. The X-axis denotes log_2_ fold-change relative to LOG, and the Y-axis represents -log_10_(adjusted *P*-value).

**Figure S3.**
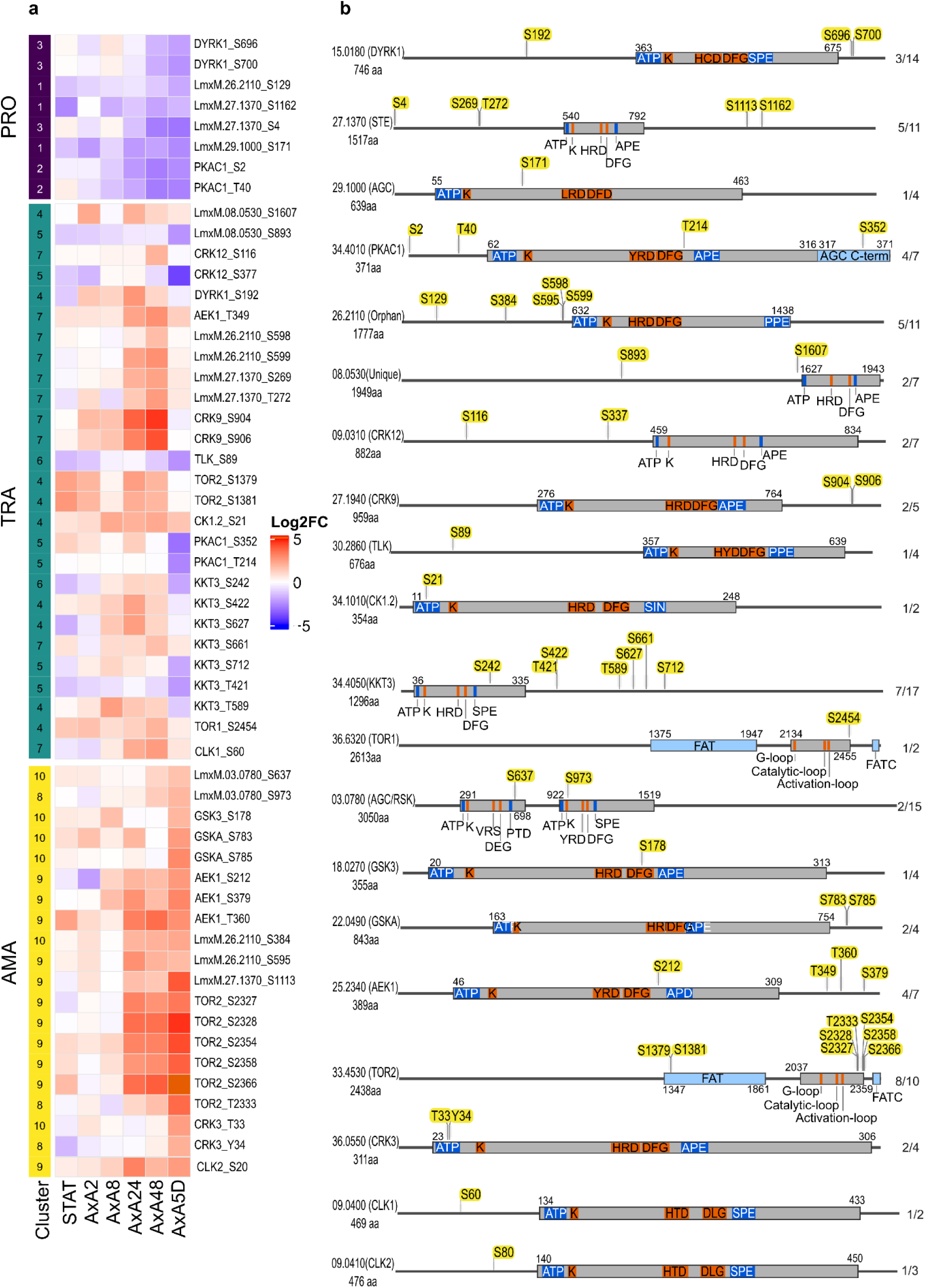
Structural architecture and phosphorylation dynamics during differentiation of protein kinases required for promastigote replication. (a) Heatmap displaying Log_2_ fold-change values of modulated phosphosites across distinct developmental stages: Promastigote stage PRO (Promastigote); TRA (Transition); AMA (Amastigote). Phosphosites were partitioned into categories derived from Mfuzz clustering. (b) Schematic representation of the 20 identified protein kinases with modulated phosphorylation sites (indicated by yellow markers). Core kinase domains are depicted as purple bars, indicating key catalytic landmarks: the ATP-binding site (ATP), the catalytic lysine (K), the catalytic loop aspartate within the HRD motif and its variants (YRD, HTD, HCD, HGD, HQD, HLD, HSD), the metal-coordination aspartate within the DFG motif and its variants (DLG, DFA, DFN, DMG, DWG) and the activation loop anchor APE motif and its variants (GPE, SPE, PPE). Additional structural and regulatory domains are represented by light blue bars, including Forkhead-associated (FHA), Phox (PX), AGC kinases C-term regulatory (AGC kin Ct), calcium binding EF hand (EF-Hand), Cytoskeleton Associated Protein Glycine-rich (CAP-Gly), FAT/FATC regulatory and PDZ scaffolding domains. Domain boundaries were identified using the PROSITE database. Core catalytic motifs were defined using multiple sequence alignment with MAFFT-SparseCore G-INS-i global iterative method. Sequence maps were generated using SnapGene software (Dotmatics). Gene IDs have been shortened for clarity (eg LmxM.07.069 to 07.0690).

**Figure S4.**
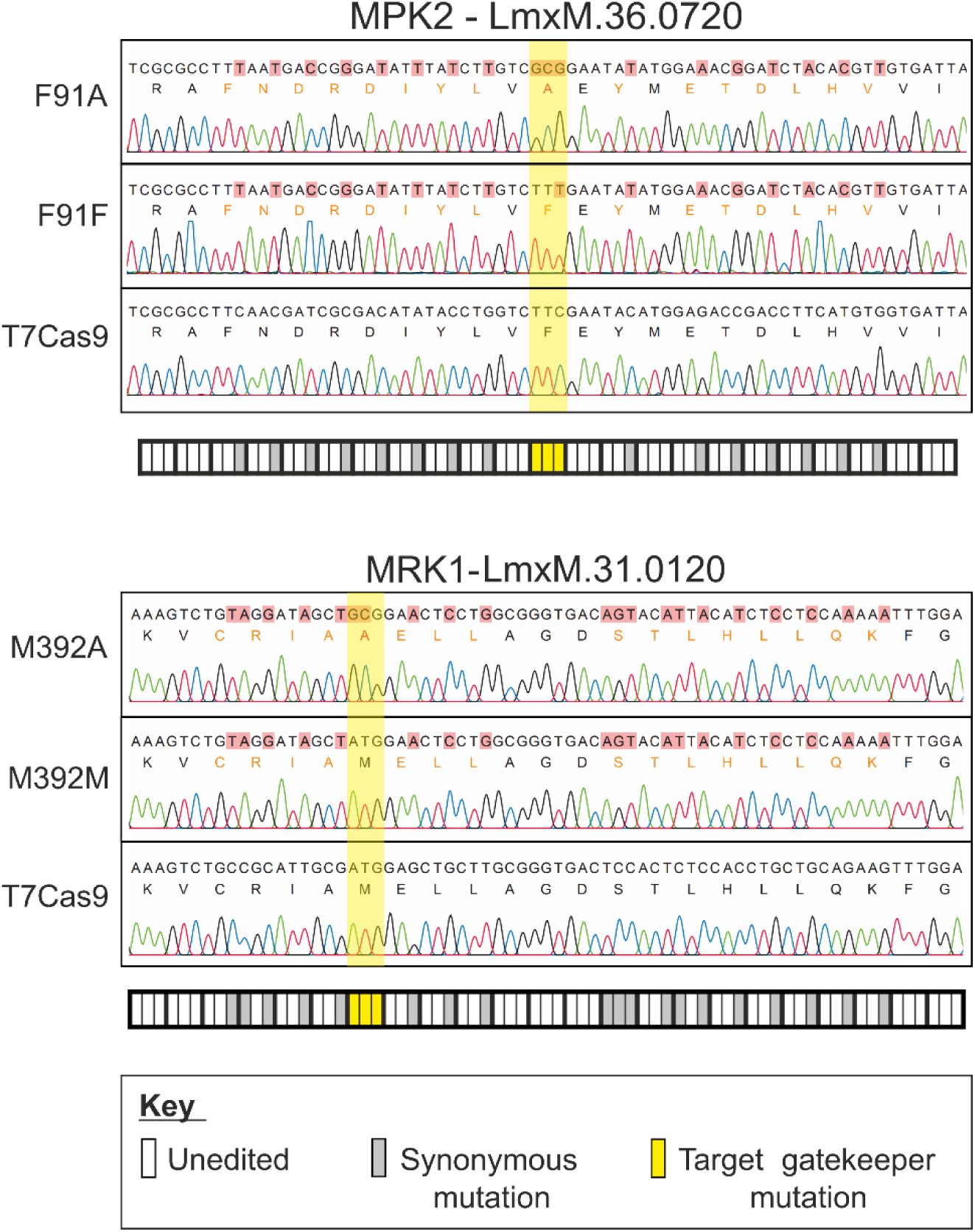
Precision editing of MPK2 and MRK1. Sanger sequencing alignments of the edited region of MPK2^F91A^, MPK2^F91F^, MRK1^M392A^ and MRK1^M392M^ mutants, or the parental cell line (T7Cas9) for the same region. Target mutation is highlighted in yellow across all cell lines. Mutated bases are highlighted with red in the aligned sequence, but are also represented below each alignment, with grey rectangles indicating synonymously recoded bases, yellow indicating nonsynonymous mutated bases, and white rectangles indicating native sequence. Triplet codons are highlighted with a thicker outline for clarity. Protein sequence for the respective region is shown below the DNA sequence of each cell line, with black text indicating the same DNA and protein sequence as the parental; orange text indicating the same protein sequence but altered DNA sequence; and red text indicating altered DNA and protein sequence compared to the parental.

**Figure S5.**
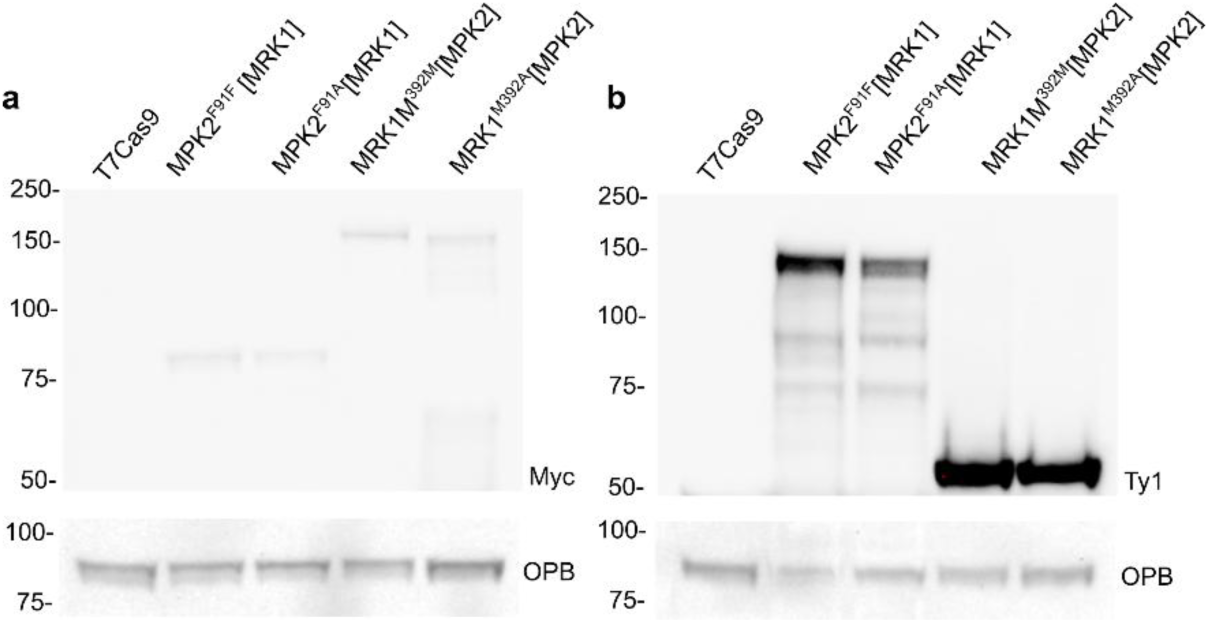
Validation of MPK2 and MRK1 silent and analogue-sensitive lines cross-complemented with the reciprocal kinase, by western blot. Myc-tagged MPK2^F91F^ (Silent) and MPK2^F91A^ (AS) lines were transfected with a plasmid overexpressing Ty1-tagged MRK1; Myc-tagged MRK1^M392M^ (Silent) and MRK1^M392A^ (AS) lines were transfected with a plasmid overexpressing Ty1-tagged MPK2. (a) Anti-Myc western blot, showing expression of the endogenously Myc-tagged kinase (MPK2 or MRK1, Silent or AS) in each background line. (b) Anti-Ty1 western blot, showing overexpression of the reciprocal, plasmid-encoded kinase in each corresponding background. Untransfected T7Cas9 parental cells were included as a negative control for Myc and Ty1 signal. 50 µg of total promastigote lysate was loaded per lane; oligopeptidase B (OPB) was used as a loading control.

**Figure S6.**
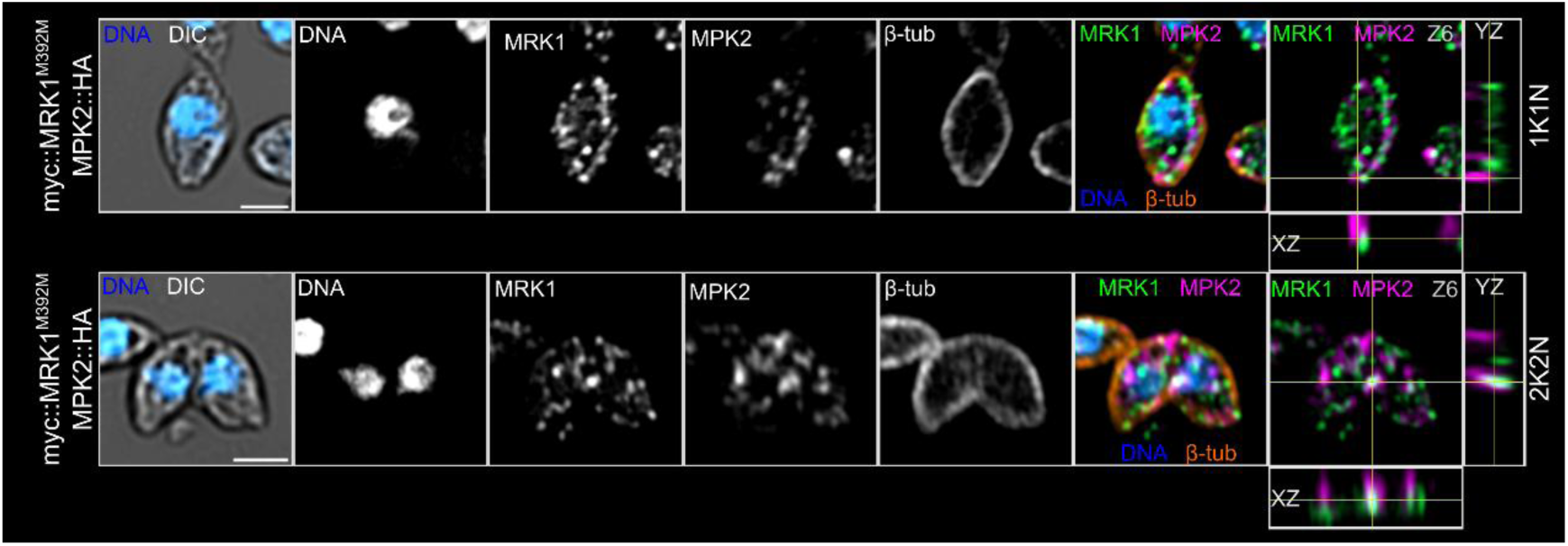
High resolution confocal immunofluorescence micrographs displaying the spatial organisation of HA::MPK2 in the myc::MRK1^M392M^ background; in mature amastigotes after 12 h with 15 µM of 1NA-PP1 treatment. Representative micrographs representing 1K1N (top) and 2K2N (bottom) cells are shown. Merged on the specified Z-stack and orthogonal views of the full Z-stack (xz and yz cross-sections at the yellow crosshair) are shown. Protein kinases were labelled with anti-myc (green) and anti-HA (magenta), microtubules were stained with KMX-1 (orange), and DNA was counterstained with DAPI (blue). Scale bar, 2µm.

**Fig. S7:**
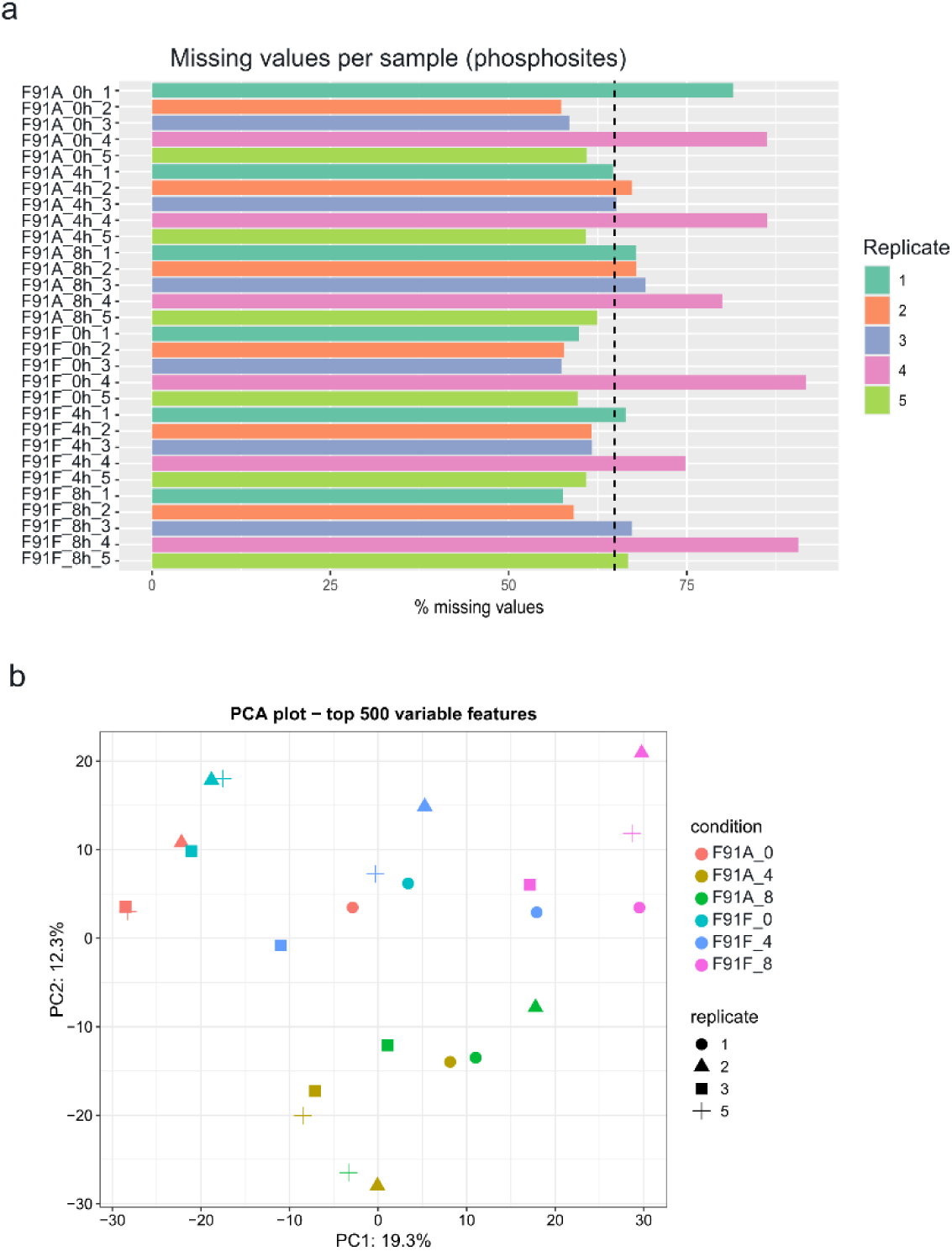
Sample quality control and imputation of the MPK2-inhibition phosphoproteome. (a) Percentages of missing values per sample in the phosphosite dataset, coloured by biological replicate. Dashed line, median across samples. Replicate 4 showed elevated missing values in all six conditions and was excluded from both the phosphosite and the whole-cell datasets. The untreated sample F91A_0h_1 was retained. (b) Principal component analysis of phosphosite intensities after vsn normalization and mixed imputation (500 most variable sites; n = 4 biological replicates per condition). PC1 separates samples by time, and PC2 separates F91A samples at 4h and 8 h, in which MPK2 is inhibited, from all F91F samples and untreated F91A (0h). F91A_0h_1 clusters with the control lines.

## Notes

### Competing Interest Statement

The authors have declared no competing interest.

